# Transcriptome-wide profiles of circular RNA and RNA binding protein interactions reveal effects on circular RNA biogenesis and cancer pathway expression

**DOI:** 10.1101/2020.03.19.997478

**Authors:** Trine Line Hauge Okholm, Shashank Sathe, Samuel S. Park, Andreas Bjerregaard Kamstrup, Asta Mannstaedt Rasmussen, Archana Shankar, Niels Fristrup, Morten Muhlig Nielsen, Søren Vang, Lars Dyrskjøt, Stefan Aigner, Christian Kroun Damgaard, Gene Yeo, Jakob Skou Pedersen

## Abstract

Circular RNAs (circRNAs) are stable, often highly expressed RNA transcripts with potential to modulate other regulatory RNAs. A few circRNAs have been shown to bind RNA binding proteins (RBPs), however, little is known about the prevalence and strength of these interactions in different biological contexts. Here, we comprehensively evaluate the interplay between circRNAs and RBPs in the ENCODE cell lines, HepG2 and K562, by profiling the expression of circRNAs in fractionated total RNA-sequencing samples and analyzing binding sites of 150 RBPs in large eCLIP data sets. We show that KHSRP binding sites are enriched in flanking introns of circRNAs in both HepG2 and K562 cells, and that KHSRP depletion affects circRNA biogenesis. Additionally, we show that exons forming circRNAs are generally enriched with RBP binding sites compared to non-circularizing exons. To detect individual circRNAs with regulatory potency, we computationally identify circRNAs that are highly covered by RBP binding sites and experimentally validate circRNA-RBP interactions by RNA immunoprecipitations. We characterize circCDYL, a highly expressed circRNA with clinical and functional implications in bladder cancer, which is covered with GRWD1 binding sites. We confirm that circCDYL binds GRWD1 *in vivo* and functionally characterizes the effect of circCDYL-GRWD1 interactions on target genes in HepG2. Furthermore, we confirm interactions between circCDYL and RBPs in bladder cancer cells and demonstrate that circCDYL depletion affects hallmarks of cancer and perturbs the expression of key cancer genes, e.g. *TP53* and *MYC*. Finally, we show that elevated levels of highly RBP-covered circRNAs, including circCDYL, are associated with overall survival of bladder cancer patients. Our study demonstrates transcriptome-wide and cell-type-specific circRNA-RBP interactions that could play important regulatory roles in tumorigenesis.

## Introduction

Circular RNAs (circRNAs) are covalently closed RNA molecules often derived from precursor mRNA (pre-mRNA) through a backsplicing event, in which a downstream 5’ splice donor backsplices to an upstream 3’ splice acceptor (1). First identified in the early 1990s, eukaryotic circRNAs were thought to be rare and a result of erroneous splicing events (2). Twenty years later, the advent of high-throughput sequencing of non-polyadenylated transcriptomes and bioinformatic analyses have made it possible to detect thousands of circRNAs, many of which are highly abundant and conserved across species (3–5). Accumulating evidence links circRNAs to development and progression of different diseases (reviewed in (6)) and several recent studies have shown that circRNAs are involved in tumorigenesis (7, 8). Due to their structural stability (4), tissue specificity (3), and relatively high expression levels in exosomes (9), blood (10), and plasma (11), circRNAs have been suggested as a new class of biomarkers and potential therapeutic targets.

RNA binding proteins (RBPs) are proteins that bind to double or single-stranded RNA. Some RBPs contain well-established RNA binding domains, including the RNA recognition motif (RRM) or the K-homology (KH) domain and bind to well-defined motifs. However, many RBPs rely on contextual features as well, e.g. secondary structure, flanking nucleotide composition, or short non-sequential motifs, complicating RNA target predictions from sequence alone (12). RBPs play crucial roles in all aspects of RNA biology, e.g. RNA transcription, pre-mRNA splicing and polyadenylation as well as modification, stability, localization, and translation of RNA (reviewed in (13)). Recent studies have shown that RBPs also affect all phases of the circRNA lifecycle (reviewed in (14)). Some RBPs are involved in circRNA biogenesis as has been shown for Quaking (QKI) (15), FUS (16), HNRNPL (17), RBM20 (18), and Muscleblind (19), which bind to specific intronic RBP motifs and promote formation of some circRNAs in certain biological settings. Besides RBP binding motifs, complementary sequences in both flanking introns, like *Alu* elements, facilitate circRNA production by RNA pairing (4). The RBP immune factors NF90/NF11 promote circRNA formation by directly binding to inverted repeated *Alus* (IR*Alus*) (20), while ADAR1 (21) and DHX9 (22) reduce circularization by destabilizing IR*Alu*-mediated RNA pairing.

The functional role of most circRNAs is still unknown. A number of circRNAs, e.g. ciRS-7 and circSRY, have been reported to function as miRNA sponges by binding a large number of microRNAs (miRNAs) and thereby regulating miRNA target genes (3, 23). More recent evidence using a ciRS-7 knockout mouse, suggests that ciRS-7 is important for normal brain function and for maintaining proper miR-7 levels (24), which in the absence of ciRS-7, becomes efficiently cleared by a long non-coding RNA (lncRNA) Cyrano (25). Other specific circRNAs have been shown to modulate host gene expression by interacting with RBPs, such as circMbl, which regulates the expression of its parent gene in a negative feedback loop between MBL and circMbl production (19). circRNAs may also regulate translation efficiency, as reported for the PABPN1 mRNA, whose translation is inhibited by the encoded circPABPN1 that efficiently “sponges” a translation stimulator RBP, HuR. (26). Additionally, circFoxo3 was found to interact with CDK2 and p21 to repress cell cycle progression in cancer cell lines (27) and to promote cardiac senescence by interacting with ID-1, E2F1, FAK, and HIF1*α* in the mammalian heart (28). Another study showed that a specific RBP, IGF2BP3, associates with several circRNAs (29).

Despite few examples of circRNA-RBP interplay, little is known about the overall ability of circRNAs to interact with RBPs. Based on binding sequence motifs of 38 RBPs and nucleotide sequence alone, You et al. found that neuronal circRNAs are not enriched with RBP binding sites compared to mRNAs (30). However, comprehensive understanding of circRNA-RBP interactions on a global scale is missing. Through extensive analysis of high throughput data sets of experimentally defined RBP binding sites combined with circRNA profiling, we can screen the entire genome for circRNA-RBP interactions and study regulatory dependencies. Since RBPs are essential to maintain normal function of the cells, defects in the expression or localization of RBPs can cause diseases (31). The abundance, high stability, and general lack of protein translation, which normally would displace most bound RBPs, make circRNAs ideal binding platforms for more than transient RBP interactions. Thus, binding and deregulation of RBPs through circRNA-RBP interactions could likely have long-term cellular effects.

Here, we evaluate the overall potential of circRNAs to interact with RBPs. Based on eCLIP data profiling the binding sites of 150 RBPs in HepG2 and K562 (32) and deep total RNA-sequencing (RNA-Seq) samples allowing circRNA quantification (33), we comprehensively study the ability of circRNAs to interact with RBPs as well as the capability of RBPs to influence circRNA formation (Figure 1A+B). We show that KHSRP binding sites are enriched in intronic regions flanking circRNAs compared to non-circularizing exons, and that KHSRP depletion diminishes circRNA expression. Additionally, we find that circularizing exons are enriched with RBP binding sites compared to non-circularizing exons indicating regulatory potency of circRNA-RBP interactions. We investigate the potential of individual circRNAs to function as RBP sponges and show experimentally that circRNAs interact with RBPs in a cell-type-specific manner. Specifically, we demonstrate that the highly expressed circCDYL is almost completely covered with GRWD1 binding sites in HepG2, and that circCDYL depletion in HepG2 cells counteracts the effect of GRWD1 depletion on target genes. In bladder cancer cell lines, we show that circCDYL interacts with IGF2BP1 and IGF2BP2, and that depletion of circCDYL or these RBPs disturb several hallmarks of cancer. Specifically, we show that some key tumor genes, including *TP53* and *MYC,* are affected by circCDYL knockdown. Finally, we show that the expression of circRNAs highly covered with RBP binding sites, including circCDYL, is positively correlated with overall survival of bladder cancer patients.

**Figure 1:**
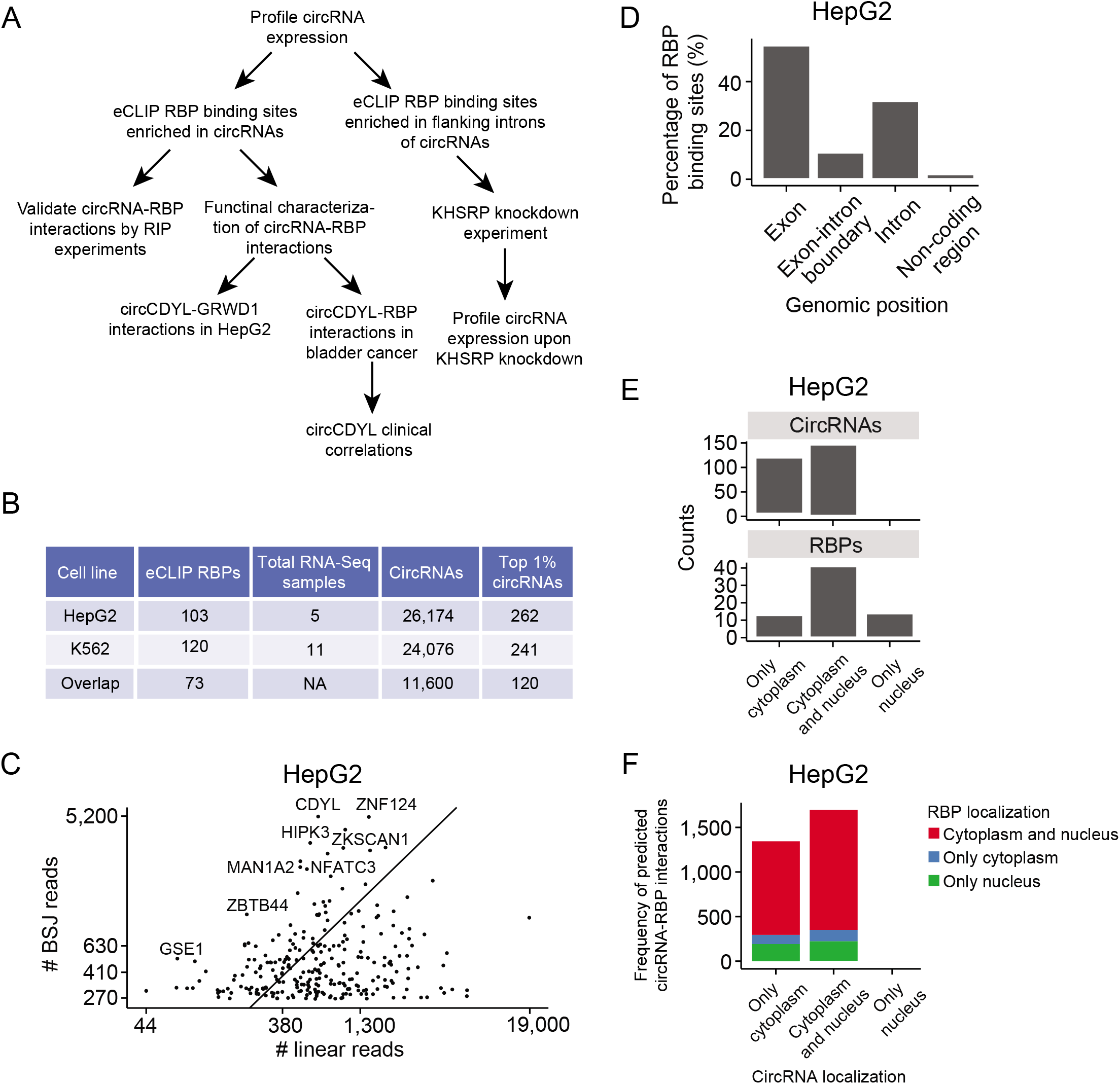
Circular RNAs are highly expressed in HepG2 and K562 and generally colocalizes with RBPs. **1A)** Flowchart overview of analyses. The initial analyses and characterizations are based on the ENCODE cell lines HepG2 and K562. Follow-up experiments are performed in HepG2 cells and bladder cancer cell lines to evaluate the relevance of findings in tumorigenesis. **1B)** Summary of input data sets and inferred sets of circRNAs. The sets of high-confidence, highly expressed circRNAs (top 1% circRNAs) are used in most of the downstream analyses. **1C)** Number of backsplice junction (BSJ) reads supporting the 1% highest expressed circRNAs (n = 262) and the number of reads spanning canonical splice sites in the corresponding linear transcripts in HepG2. X-and Y-axis are plotted on a logarithmic scale (log10) showing actual counts. Some circRNAs are depicted with their host gene name. **1D)** Genomic location of RBP binding sites in circRNA loci in HepG2. Most RBP binding sites are found within exonic parts of the circRNAs. Exonic parts span 0.185 Mb, while introns span 2.706 Mb. **1E)** Localization of circRNAs (n = 262; top) and RBPs (n = 68; bottom) across cellular compartments of HepG2. For circRNAs, all (except for one) are found in the cytoplasm (n = 261), with a large subset that is also found in the nucleus (n = 146). No circRNAs are exclusively expressed in the nucleus. For RBPs, most are located in both cellular compartments (n = 41), while some RBPs are expressed exclusively in the cytoplasm (n = 13) or the nucleus (n = 14). **1F)** Localization of circRNAs and RBPs that are predicted to interact from eCLIP data in HepG2. CircRNA localization is shown on x-axis, while RBP localization is indicated by color. For 94% of the predicted circRNA-RBP interactions, the circRNA and RBP are expressed in the same subcellular compartment.

## Results

### Circular RNAs are highly expressed in HepG2 and K562 and generally colocalizes with RBPs

To identify and quantify the expression of circRNAs in HepG2 and K562 cells, we utilized all available ENCODE whole transcriptome RNA-Seq data sets for these cell lines (HepG2; n = 5 and K562; n = 11) (Supp. file 1+2, Supp. Table 1) (33). Employing the CIRI2 pipeline (34), we detected 26,174 and 24,076 unique circRNAs supported by at least two reads spanning the backsplice junction (BSJ) in HepG2 and K562, respectively, of which 11,600 circRNAs were identified in both cell lines (Figure 1B, Supp. Table 2+3). Of all identified circRNAs, 44% (HepG2) and 43% (K562) are described in circBase, a large compendium of circRNAs from different studies (35). As many circRNAs are lowly expressed and hard to distinguish from artifacts (36) (Supp. Figure 1A), we generated two sets of high-confidence circRNAs by considering only the 1% highest expressed circRNAs in each cell line (Figure 1B). These sets contain 262 circRNAs in HepG2, each supported by at least 260 reads, and 241 circRNAs in K562, each supported by at least 210 reads. The vast majority of these highly expressed circRNAs are found in circBase (94% in HepG2 and 91% in K562). We compared the expression of the circRNAs to their corresponding linear transcript and found that many circRNAs (39%) in HepG2 are more expressed than the linear counterpart, among these common circles like circHIPK3 and circCDYL (Figure 1C (HepG2), Supp. Figure 1B (K562)). Corroborating previous findings, we observed that most circRNAs are comprised of 5 or less exons (3) (Supp. Figure 1C (HepG2 and K562)), and that exons giving rise to 1-exon-circRNAs are significantly longer than exons giving rise to multiple-exon-circRNAs (4) (P < 2.2e-10, Wilcoxon Rank Sum Test, Supp. Figure 1D (HepG2 and K562)).

Generally, circRNAs are only comprised of exonic sequences as introns are spliced out in the circularization process (4). Based on eCLIP data and a high-confidence set of peak calls representing binding sites of RBPs in HepG2 (n = 103) and K562 (n = 120) (Methods, Supp. files 3+4), we evaluated the genomic positions of RBP targets encoded by the circRNA loci (Supp. Table 4). 55% of the RBP binding sites are within exonic parts of the circRNAs (span: 0.185 Mb), while 32% of the binding sites are found strictly in introns (span: 2.706 Mb) and 11% at exon-intron boundaries in HepG2 (Figure 1D, Supp. Figure 1E (K562)). The remaining 2% are found in non-coding regions within circRNAs. As introns are usually spliced out and RBPs that bind to exon-intron boundaries are likely involved in normal pre-mRNA splicing (37), we focused initially on RBPs with binding sites in exonic regions of circRNAs. To be able to interact, circRNAs and RBPs should be present in the same cellular compartments of the cell. Based on ENCODE subcellular fractionated RNA-Seq expression data and immunofluorescence imaging of RBP occupancy in HepG2 (38), we evaluated the localizations of circRNAs and RBPs in HepG2. We found that all circRNAs are located in the cytoplasm as ordinarily the case (4, 5, 39) (Figure 1E). Though some circRNAs are also found in the nucleus, no circRNAs are more highly expressed in the nucleus than in the cytoplasm (Supp. Figure 1F). For the RBPs (n = 68), most are found in both compartments (n = 41), while some are found exclusively in the cytoplasm (n = 13) and some in the nucleus (n = 14) (Figure 1E, Supp. Table 5). For 94% of the circRNA-RBP interactions inferred from eCLIP data, we found that the involved circRNA and RBP were co-located in the same subcellular compartments, indicating potential to interact (Figure 1F).

### KHSRP binding is enriched in introns flanking circRNAs and affects biogenesis

To comprehensively analyze if RBPs influence circRNA formation in HepG2 and K562 cells, we evaluated whether binding sites for individual RBPs are enriched in introns flanking circRNAs compared to non-circularizing exons. We divided the genomic regions into different sets (Figure 2A): 1) the subset of highly expressed, high-confidence circRNAs; 2) all other circRNAs; and 3) non-circularizing exons (non-circ-exons) from genes producing circRNAs. For the latter, we excluded the first and last exons in each gene as they are not surrounded by introns on both sides.

**Figure 2:**
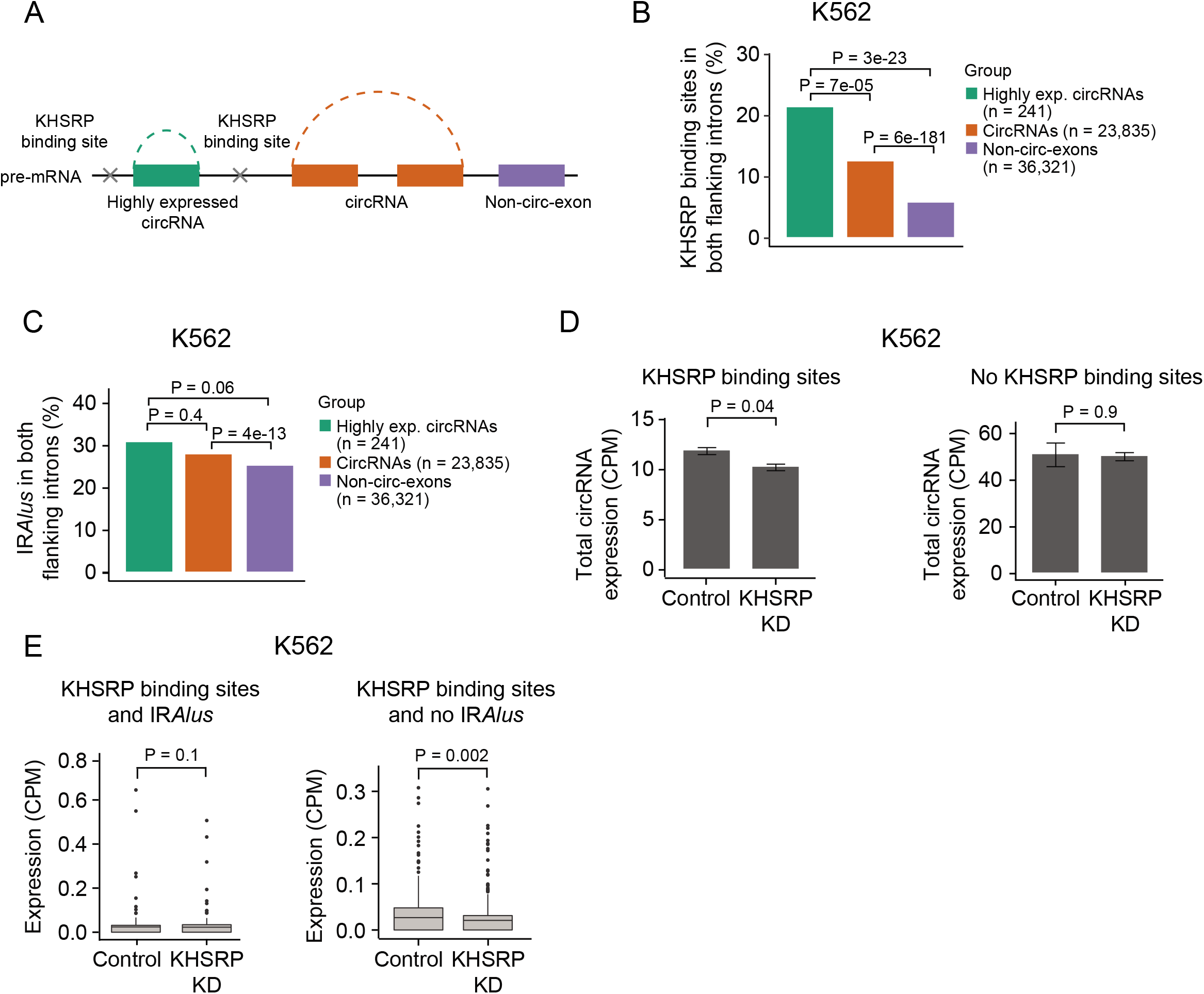
KHSRP binding is enriched in introns flanking circRNAs and affects biogenesis. **2A)** Illustration of circRNA and exon categories and intronic location of KHSRP binding sites. The categories are defined as the top 1% highest expressed circRNAs (green), all other circRNAs (orange), and non-circularizing exons from genes producing circRNAs (purple). First and last exons in each gene are disregarded for the analyses. **2B)** Percentage of circRNAs and non-circ-exons in genes producing circRNAs with KHSRP binding sites in both flanking introns in K562. P-values obtained by Chi-square Test. **2C)** Percentage of circRNAs and non-circ-exons in genes producing circRNAs with inverted repeated *Alu* elements (IR*Alus*) in both flanking introns in K562. P-values obtained by Chi-square Test. **2D)** Total expression of circRNAs with (left, n = 297) and without (right, n = 1,537) KHSRP binding sites in both flanking introns in KHSRP knockdown (KD) and control samples in K562. P-values obtained by T-test. CPM = Counts per million. **2E)** Expression of circRNAs with KHSRP binding sites and with (left, n = 95) or without (right, n = 202) IR*Alus* in both flanking introns in KHSRP knockdown (KD) and control samples in K562. P-values obtained by T-test.

For all individual RBPs, we evaluated the presence of binding sites in both intronic regions (10 kb to each side) of circRNAs and non-circ-exons (Supp. Figure 2A+B (K562 and HepG2)). Noteworthy, we found that > 20% of the highly expressed circRNAs (n = 241) and > 10% of all other circRNAs (n = 23,835) in K562 possess KHSRP binding sites in both flanking introns, which is 3.6 and 2.1 times more than non-circ-exons (n = 36,321), respectively (all P < 0.001, Chi-square Test, Figure 2B). Similar significant results were observed in HepG2, although fewer circRNAs are surrounded by KHSRP binding sites (all P < 0.001, Chi-square Test, Supp. Figure 2C).

Since IR*Alus* can also facilitate circularization, we evaluated the existence of IR*Alus* in both flanking introns of circRNAs and non-circ-exons. IR*Alus* are only weakly enriched for both highly expressed circRNAs (1.2x, P = 0.06, Chi-square Test) and all other circRNAs (1.1x, P = 4e-13, Chi-square Test) compared to non-circ-exons in K562 (Figure 2C). The same pattern is observed for HepG2 cells (Supp. Figure 2D). This indicates that IR*Alus* are not a major driver of circularization in K562 and HepG2 cells. Specifically, flanking introns of circRNAs with KHSRP binding sites were not enriched with IR*Alus* (Supp. Figure 2E (K562 and HepG2)).

KHSRP binds to single-stranded RNA and exerts diverse functions in RNA metabolism, e.g. by promoting mRNA decay, inducing miRNA maturation, and affecting alternative RNA splicing (40, 41). Based on immunofluorescence imaging in HepG2, we see that KHSRP is only detectable in the nuclear fraction of the cell (Supp. Table 5). We hypothesize that KHSRP binds to flanking introns of circRNAs in the pre-mRNA transcript in the nucleus and thereby promotes circRNA biogenesis.

To test our hypothesis, we profiled the expression of circRNAs using CIRI2 upon KHSRP knockdown (KD) in K562 and HepG2 cells (Methods). Overall, we found no difference in total circRNA expression between K562 control and KHSRP KD samples (n = 1,834) (P = 0.6, T-test, Supp. Figure 2F). However, for circRNAs with KHSRP binding sites in both flanking introns (KHSRP-circRNAs, n = 297), we found a 15% decrease in circRNA expression upon KHSRP KD (P = 0.04, T-test, Figure 2D), while there was no effect on expression levels for circRNAs without KHSRP binding sites in both flanking introns in K562 (n = 1,537) (P = 0.9, T-test, Figure 2D). For KHSRP-circRNAs, we evaluated the presence of IR*Alus* in both flanking introns. We found that the expression of KHSRP-circRNAs without IR*Alus* in flanking introns (n = 202) is more affected by KHSRP depletion (P = 0.002, Wilcoxon Rank Sum Test, Figure 2E) than KHSRP-circRNAs surrounded by IR*Alus* (n = 95, P = 0.1, Wilcoxon Rank Sum Test, Figure 2E), supporting the role of KHSRP in the biogenesis of a subset of circRNAs.

Since our observations could be explained by overall splicing perturbations in KHSRP KD samples, we evaluated the expression of the corresponding linear RNA of circRNAs. We found no difference in linear RNA expression between KHSRP KD and control samples for circRNAs with (P = 0.9, T-test, Supp. Figure 2G) or without KHSRP binding sites (P = 0.5, T-test, Supp. Figure 2G), indicating that KHSRP specifically affect the expression of circRNAs.

We observed no effect of KHSRP KD on circRNA expression levels of circRNAs with (P = 0.5, T-test, Supp. Figure 2H) or without (P = 0.1, T-test, Supp. Figure 2H) KHSRP binding sites in both flanking introns in HepG2. This could be explained by lower expression levels of KHSRP and a generally lower fraction of circRNAs with KHSRP binding sites in flanking introns (Supp. Figure 2C).

Taken together, our results identify KHSRP as an RBP that appear to be involved in the biogenesis of a subset of circRNAs with KHSRP binding sites in flanking introns in K562 cells.

### Exons comprising circRNAs are enriched with RBP binding sites

Most circRNAs consist of protein coding exons. If RBP binding is enriched in circRNAs, it suggests a regulatory layer in addition to their protein-coding capacity.

To evaluate the enrichment of RBP binding sites in circRNAs, several considerations had to be taken into account. Since circRNAs share sequence with their cognate linear transcript, RBP targets from eCLIP data do not directly distinguish between interactions with circular or linear RNA transcripts. Additionally, gene expression levels influence the ability to detect RNA-RBP interactions. By dividing all genes into expression deciles, we found that the fraction of exons covered with RBP binding sites (RBP-coverage) increases with transcript abundance as expected (r > 0.4, P < 2.2e-16, Pearson’s product-moment correlation, Supp Figure 3A (HepG2 + K562)).

To evaluate whether circRNAs show more RBP enrichment than expected simply from the expression level of their parent gene, we compared the RBP-coverage of exons comprising circRNAs to linear exons that are not involved in circularization within different comparable gene sets: A) circRNA host genes; B) genes of the same expression level; and C) the most highly expressed genes. We divided exons into three different groups: 1) backsplice junction circularizing exons (BSJ circ-exons), which are directly involved in backsplicing; 2) circularizing exons (circ-exons), that are potentially part of circRNAs but not supported by BSJ reads; 3) and non-circ-exons, which are not part of circRNAs (Figure 3A). Since many RBPs regulate transcription and translation by binding to the 5’-and 3’-UTRs of mRNAs (42) and since they give rise to few circRNAs, we disregarded the first and last exons in each gene (Supp. Figure 3B (HepG2 and K562)).

**Figure 3:**
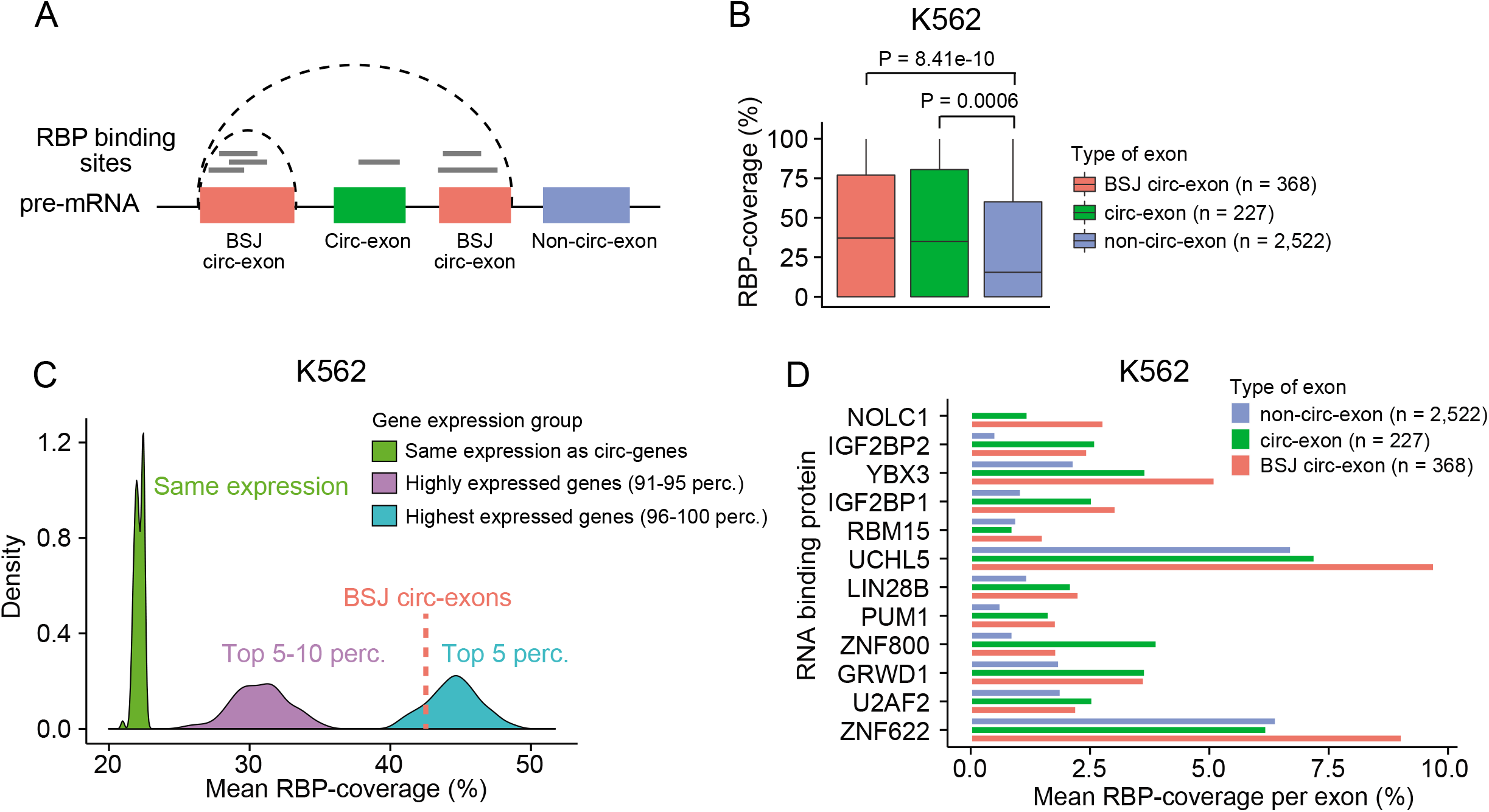
Exons comprising circRNAs are enriched with RBP binding sites. **3A)** Illustration of exon categories in genes forming circRNAs. Backsplice junction circularizing exons (BSJ circ-exons, red) are involved in the backsplicing event. Circularizing exons (circ-exons, green) are potentially part of circRNAs, if not spliced out. Non-circularizing exons (non-circ-exons, blue) are not part of circRNAs. Grey lines represent RBP binding sites. To evaluate the fraction of exons covered by RBP binding sites overall, the individual overlapping RBP binding sites were merged. **3B)** RBP binding site coverage for each category of exons in genes that produce highly expressed (top 1%) circRNAs in K562 (n = 185). P-values obtained by Wilcoxon Rank Sum Test. **3C)** Comparison of RBP-coverage between BSJ circ-exons and exons in groups of genes of different expression levels (K562). The mean RBP-coverage of the highly expressed BSJ circ-exons (n = 368) is 43% (red punctuated line; median = 37%). Exons randomly sampled from genes while ensuring the same expression profile as genes producing highly expressed circRNAs (circ-genes) have a much lower mean coverage of 22% (green; median = 0%). Exons of highly expressed genes (top 5-10 percentiles) also showed a lower mean coverage of 31% (purple; median = 6%), while the most highly expressed genes (top 5 percentiles) had a slightly higher mean of 45% (blue; median = 42%). The random sampling procedures were repeated with 100 iterations. Empirical P-values for: BSJ circ-exons vs. same expression, P < 0.01; BSJ circ-exons vs. Top5-10, P < 0.01; BSJ circ-exons vs Top5, P = 0.83. **3D)** Mean RBP-coverage per exon for individual RBPs (K562). All RBPs shown here have significantly more target sites in BSJ circ-exons than in non-circ-exons of the same genes (FDR < 0.1, Wilcoxon Rank Sum Test). Only RBPs with at least 20 distinct binding sites in total are considered.

Interestingly, for the high-confidence, highly expressed circRNAs, we found that BSJ circ-exons as well as circ-exons were significantly more covered with RBP binding sites than internal non-circ-exons within the same genes for both K562 (all P < 0.001, Wilcoxon Rank Sum Test, Figure 3B) and HepG2 (all P <= 0.05, Wilcoxon Rank Sum Test, Supp. Figure 3C). The difference was also significant when examining the complete set of circRNAs and their associated non-circ-exons (all P < 2.2e-16, Wilcoxon Rank Sum Test, Supp. Figure 3D (HepG2 + K562)).

To compare RBP-coverage of BSJ circ-exons to genes of the same expression levels as genes producing highly expressed circRNAs (circ-genes), we divided all genes into expression level percentiles. We randomly drew non-circ-exons, while ensuring the same expression level distribution as circ-genes, which are distributed across percentiles ~20-100 (Supp. Fig 3E (HepG2 + K562)). We found that BSJ circ-exons are significantly more covered with RBP binding sites than non-circ-exons of the same expression level in both cell lines (P < 0.01, Empirical p-value, Figure 3C (K562), Supp Figure 3F (HepG2)).

Even when BSJ circ-exons were compared against the highest expressed genes, their RBP-coverage were significantly higher (P < 0.01, percentiles 90-95) or similar (P not significant, percentiles 96-100), indicating specific circRNA-RBP interactions (Figure 3C (K562), Supp. Figure 3F (HepG2)).

We evaluated whether any individual RBPs preferentially bind to circRNA exons. Several RBPs have significantly higher RBP-coverage on average in BSJ circ-exons compared to non-circ-exons in circ-genes (false discovery rate (FDR) < 0.1 for all shown RBPs, Wilcoxon Rank Sum Test, Figure 3D (K562), Supp. Figure 3G (HepG2)). We found that some RBPs are more prone to bind to BSJ circ-exons in both cell lines, e.g. GRWD1 and UCHL5, while others seem to be cell line specific, like NOLC1, IGF2BP1, and YBX3 in K562. We performed GO enrichment analysis to identify GO terms that are overrepresented in the subset of RBPs that bind more to BSJ circ-exons (K562; n = 12 and HepG2; n = 8). Although, we observed no significant results, we found that a larger fraction of RBPs that are more prone to bind to BSJ circ-exons are associated with mRNA stability (3.1 fold), mRNA transportation (1.8 fold), and binding to the mRNA 5’-UTR (8.3 fold) and 3’-UTR (2.8 fold) compared to the rest of the RBPs in K562 (Supp. Table 6). These results suggest that circRNAs could influence transcript expression and function by interacting with RBPs that are involved in mRNA maturation and localization. It is also possible that these RBPs facilitate circRNA transportation in a manner similar to their functions on mRNA transcripts.

Finally, we explored the ability of RBPs to interact with the unique BSJ of circRNAs. For this, we aligned eCLIP reads to a constructed reference set of all possible backsplicing events based on annotated splice sites (Methods, Supp. Figure 3H). Some potential backsplicing events were covered by eCLIP reads in HepG2 (n = 111) and K562 (n = 133) (Supp. Figure 3I (HepG2 and K562)), however, none of these correspond to a BSJ of the circRNAs we called.

Overall, our results show that exons comprising circRNAs are enriched with RBP binding sites and that some RBPs preferentially bind to circRNAs. Though RBP binding sites overlapping BSJ circ-exons could simply stem from binding to the linear form of the exons, circularizing exons generally had much higher RBP-coverage than comparable linear exons, supporting that they specifically interact with RBPs.

### CircRNAs interact with RBPs in a cell-type-specific manner

Next, we wanted to identify specific circRNAs enriched with RBP binding sites that could potentially function as RBP sponges or in other ways interact with RBPs to deregulate the expression, function, or localization of RBPs. We evaluated the coverage of RBP binding sites in the exonic parts of circRNAs (Supp. Table 7). Many known and highly expressed circRNAs are completely or almost completely covered by RBP binding sites in one or both of the cell lines, e.g. circRNAs arising from the genes *RBM39, GSE1, SMARCA5, RBM33, ZKSCAN1*, and *CDYL* (Figure 4A (HepG2 left and K562 right)). Overall, RBP-coverage is not significantly associated with the number of exons comprising the circRNAs (Supp. Figure A (HepG2 and K562)). We compared the RBP-coverage in circRNAs to the RBP-coverage of internal non-circularizing exons transcriptome-wide, and found that several of the above mentioned circRNAs fall in the 5% tail (as indicated by stars) in one, e.g. circGSE1 and circSMARCA5, or both cell lines, e.g. circCDYL and circRBM33 (Figure 4B, Supp. Table 7).

**Figure 4:**
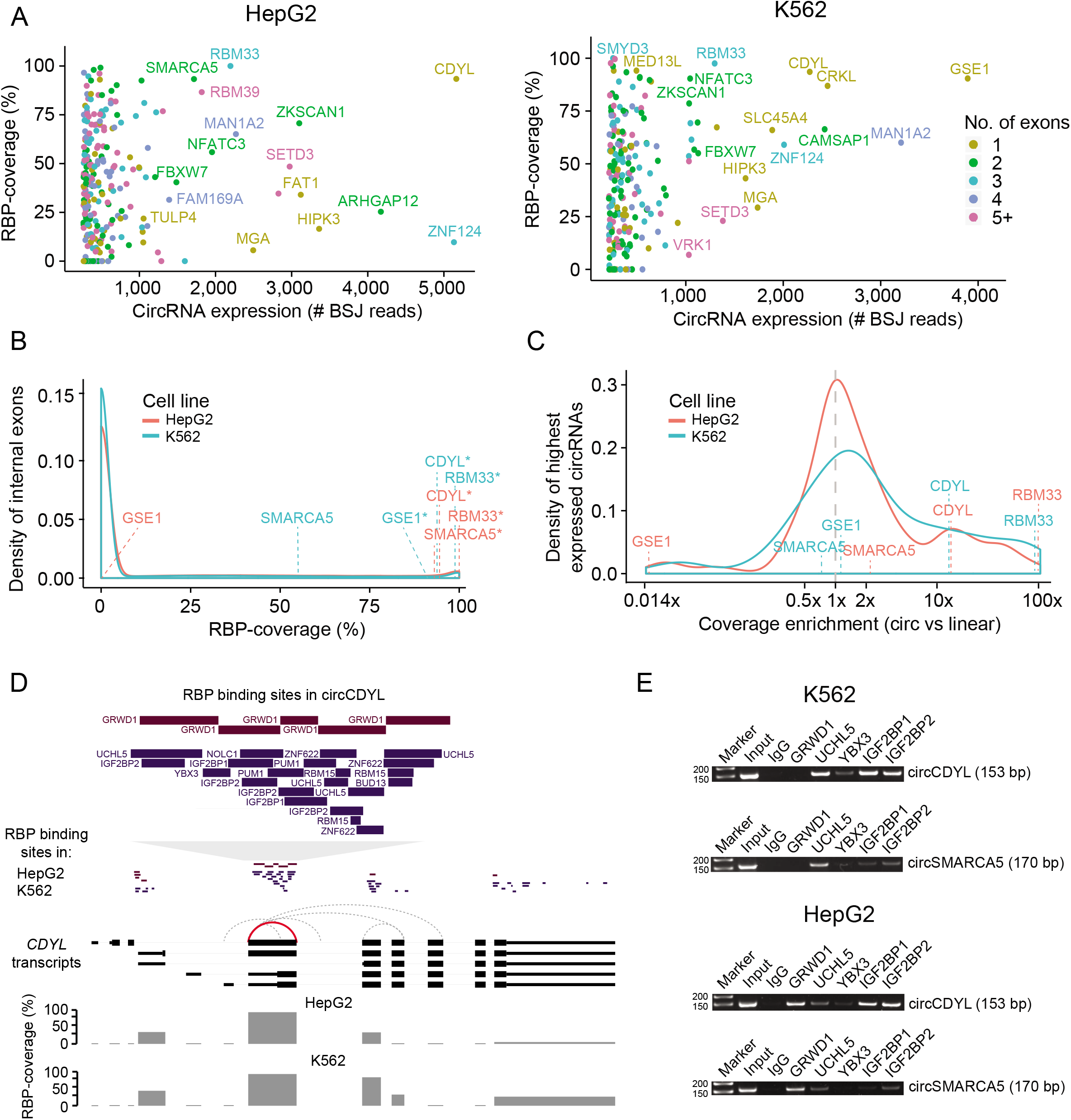
CircRNAs interact with RBPs in a cell-type-specific manner. **4A)** Percentage of the exonic part of circRNAs that is covered with RBP binding sites in HepG2 (right) and K562 (left). Only the top 1% highest expressed circRNAs are shown. Some circRNAs are depicted with their host gene name. Some circRNAs are highly covered with RBP binding sites in both cell lines, e.g. circCDYL and circRBM33. Colours indicate the number of exons constituting each circRNA. We only consider circRNAs with annotated exons. **4B)** Comparison of RBP-coverage for individual highly expressed circRNAs and all internal non-circ-exons in expressed genes in HepG2 (red) and K562 (blue). Common circRNAs that are highly covered with RBP binding sites in one or both cell lines are depicted with their host gene name. Stars indicate an RBP-coverage in the 5% tail. **4C)** Enrichment of RBP-coverage of circRNAs (top 1%) compared to the mean RBP-coverage of all non-circ-exons of their host gene in HepG2 (red) and K562 (blue). Most circRNAs contain an equal amount or slightly more RBP binding sites than non-circ-exons in their parent gene (=>1x enrichment), while some circRNAs are highly enriched with RBP binding sites, e.g. circCDYL (>10x) and circRBM33 (100x). **4D)** RBP binding sites in exons of the *CDYL* gene. Exons of the five main *CDYL* transcripts (thick part indicates protein-coding regions) and all circRNAs (stipulated lines) supported by at least two reads are shown. The highly expressed circCDYL (chr6:4891946-4892613; supported by 5,163 (HepG2) and 2,272 (K562) reads) is indicated by a red line. This exon is highly covered by RBP binding sites in both cells. Zoom-in at the top shows the binding sites and names of the individual RBPs in both cell lines. All other *CDYL*-circRNAs are supported by less than 22 reads. Bars show percentage of each exon covered with RBP binding sites. **4E)** Validation of circRNA-RBP interactions from RNA immunoprecipitation. All circRNA-RBP interactions, except for one, were verified. Here, RNA immunoprecipitation (RIP) experiments confirmed that circCDYL interacts with several RBPs in K562, e.g. UCHL5, IGF2BP1, IGF2BP2, but not with GRWD1. In HepG2, RIP experiments confirmed strong circCDYL-GRWD1 interactions. Predicted interactions between circSMARCA5 and UCHL5 in K562 and GRWD1 in HepG2 were also validated. 50 bp markers used.

To ensure our observations are not simply explained by host gene expression levels, we compared the RBP-coverage of highly expressed circRNAs to non-circularizing exons within the parent gene (Supp. Table 7). We found that most circRNAs (69%) have the same or higher RBP-coverage than non-circularizing exons in the same gene (Figure 4C (HepG2 and K562)). Of these, 20% are highly enriched (10x). Our results indicate that gene expression cannot explain the high RBP-coverage of specific circRNAs, as some circRNAs differ in RBP-coverage between cell lines despite similar circRNA and host gene expression levels. For instance, circGSE1 as well as non-circ-exons in *GSE1* are highly covered with RBP binding sites in K562 (1.2x), while there are no RBPs overlapping circGSE1 in HepG2 but many RBP binding sites in *GSE1* non-circularizing exons (ratio = 0.02). In both cell lines, circGSE1 is among the 1% highest expressed circRNAs and the parent gene *GSE1* is found in similar gene expression percentiles in K562 (81) and HepG2 (83).

Interestingly, circCDYL (11x) and circRBM33 (100x) are highly covered with RBP binding sites compared to non-circ-exons in their parent gene in both cell lines (Figure 4C). These enrichments are significantly higher than for other genes with the same number of non-circ-exons (P < 0.05, Empirical p-value). The *RBM33* gene is highly expressed in both K562 (percentile 73) and HepG2 (percentile 91). The high enrichment for circRBM33 is due to 100% RBP-coverage of circRBM33 and no RBP binding sites in the two non-circ-exons in the gene. The *CDYL* host gene, which gives rise to the highest expressed circRNA, is itself modestly expressed in both K562 (percentile 48) and HepG2 (percentile 62). CircCDYL is 93% covered with RBP binding sites in both cell lines while the non-circ-exons in *CDYL* have a mean RBP-coverage of only 7% (Figure 4D). Remarkably, the RBPs binding to the circCDYL exon differ between the cell lines. In HepG2, only one RBP, GRWD1, is binding across almost the entire sequence, while nine different RBPs have at least one binding site in K562. We observe that cell-type-specific circRNA-RBP interactions is a general phenomenon. Between 120 circRNAs that are highly expressed in both cell lines (Supp. Figure 4B) and 34 RBPs evaluated in both cell lines (Supp. Figure 4C), less than 15% of circRNA-RBP interactions (n = 1,073) are shared between the cell lines (Supp. Figure 4D).

To evaluate the validity of the predicted circRNA-RBP interactions based on the eCLIP data, we performed RNA immunoprecipitation (RIP) of five RBPs; IGF2BP1, IGF2BP2, GRWD1, YBX3, and UCHL5 that are predicted to bind to one or several of four highly expressed and highly RBP-covered circRNAs; circCDYL, circRBM33, circZKSCAN1, and circSMARCA5 (Supp. Figure 4E (HepG2 and K562)). RBP antibody specificity were validated using western blots (Supp. Figure 4F (HepG2 and K562)). We designed divergent primers against the unique backsplice junction of the circRNAs to specifically verify circRNA pull-down in RIP experiments by RT-PCR (Supp. Table 8). We evaluated all possible interactions between the RBPs and circRNAs and all, except for one, were verified. In K562, the RIP experiments confirmed that circCDYL interacts with UCHL5, YBX3, IGF2BP1, and IGF2BP2 and that both circSMARCA5, circZKSCAN1, and circRBM33 binds UCHL5 (Figure 4E, Supp. Figure 4G (K562)). In HepG2, the RIP experiments confirmed interactions between GRWD1 and all four circRNAs (Figure 4E, Supp. Figure 4G (HepG2)). There were no eCLIP data of IGF2BP2 in HepG2 but strong bands in the IGF2BP2 immunoprecipitation (IP) experiment showed that IGF2BP2 interacts with circZKSCAN1 and circCDYL. The only interaction that could not be confirmed was between GRWD1 and circZKSCAN1 in K562 (Supp. Figure 4G (K562)). One likely explanation is that GRWD1 only interacts with the linear form of *ZKSCAN1* or with circZKSCAN1 under certain conditions. Faint bands indicate that the circRNAs interact with other RBPs than the ones identified from the eCLIP data with lower affinity. The lack of low-affinity and non-specific RBP binding sites in the eCLIP data could be explained by the stringent cut-off used, of 8-fold enrichment in IP compared to input (Methods). In accordance with the eCLIP data, our results show that circRNAs interact with RBPs in a highly cell-type-specific manner.

### Functional studies of circCDYL-RBP interactions in HepG2

There are several ways in which circRNAs could interact with and regulate RBPs. CircRNAs could function as protein decoys and retain certain RBPs to specific cellular compartments, or as scaffolds to facilitate contact between two or more RBPs, or as a unit in larger functional complexes.

To understand the regulatory potential of circRNA-RBP interactions in more depth, we focused our attention on circCDYL’s interaction with RBPs. CircCDYL is one of the highest expressed circRNAs across tissues in both humans and mice (43) and is deregulated in diseases, including cancer and myotonic dystrophy (9, 44, 45). However, little is known about its regulatory functions and its potential to interact with RBPs remains unexplored.

Here, we found that circCDYL is one of the highest expressed circRNAs in both cell lines and almost entirely covered with RBP binding sites (Figure 4A (HepG2 and K562)). As seen for most circRNAs, circCDYL levels are highest in the cytosol and the circle is more abundant than its linear counterpart in both HepG2 (Figure 5A) and K562 (Supp. Figure 5A). All RBPs confirmed to interact with circCDYL are expressed in the cytoplasm as well (Supp. Table 5). Therefore, we hypothesized that circCDYL interacts with RBPs in the cytoplasm and regulates associated RBP target genes.

**Figure 5:**
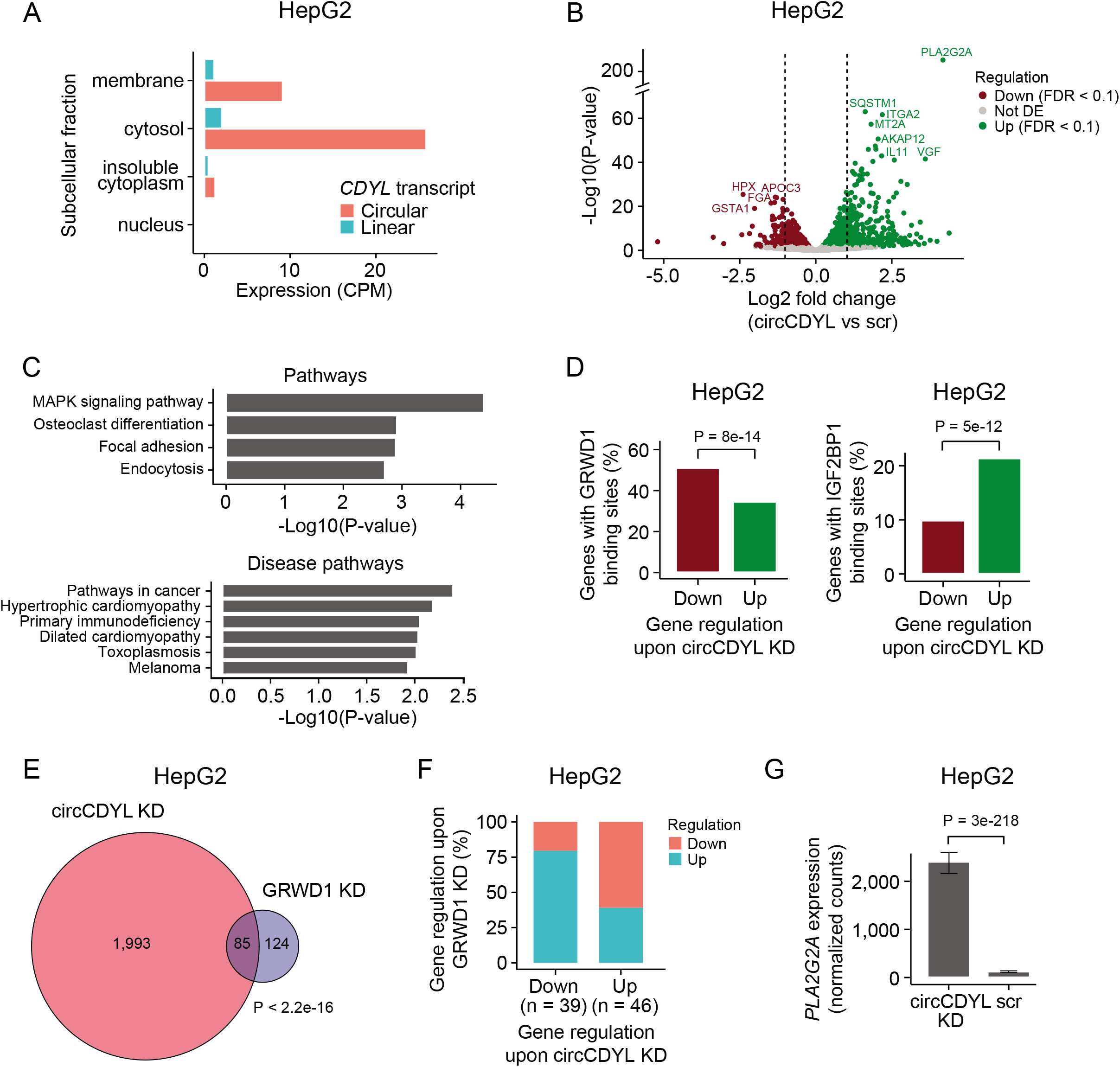
Functional studies of circCDYL-GRWD1 interactions in HepG2. **5A)** Expression of circCDYL and the corresponding linear transcript in subcellular fractions of HepG2. **5B)** Differential expression analyses of mRNAs upon circCDYL KD in HepG2. The log2 fold changes (circCDYL KD vs scr) are plotted against the negative log10(P-values). Colors indicate if genes are significantly down-(red) or upregulated (green) or not significantly differentially expressed (Not DE, grey) after Benjamini-Hochberg correction, FDR < 0.1. Vertical lines indicate a log2FC > 1 or < −1. **5C)** KEGG pathway (top) and KEGG disease pathway (bottom) analyses upon circCDYL KD in HepG2. **5D)** Percentage of significantly down-and upregulated genes upon circCDYL KD with GRWD1 (left) and IGF2BP1 (right) binding sites in HepG2. Colors indicate gene regulation upon circCDYL KD (as in Figure 5B). Only genes expressed in both data sets are considered. P-value obtained by Wald Test. **5E)** Overlap of genes affected by circCDYL KD and GRWD1 KD in HepG2. P-value obtained by Fisher’s Exact Test. **5F)** Regulation of genes affected by both circCDYL KD and GRWD1 KD in HepG2 (n = 85). The x-axis indicates gene regulation upon circCDYL KD. Percentage on y-axis and colors show how these genes are regulated upon GRWD1 KD. Downregulated genes upon circCDYL KD are mainly upregulated upon GRWD1 KD. **5G)** Expression of *PLA2G2A* upon circCDYL KD in HepG2 cells. P-value obtained by Wald Test.

To functionally characterize circCDYL, we conducted siRNA-mediated knockdown (KD) in HepG2 with an siRNA specifically recognizing the backsplice junction of circCDYL (Methods, Supp. Table 9). We validated KD efficiency by qRT-PCR (Supp. Figure 5B (HepG2), Supp. Table 10), harvested RNA in triplicates, and quantified mRNA expression using QuantSeq (46). We identified 2,233 genes that are differentially expressed (DE) upon circCDYL KD, of which 1,255 are upregulated and 978 are downregulated (Figure 5B (HepG2)). KEGG pathway analyses showed that pathways in cancer and specifically the MAPK signaling pathway are activated upon circCDYL KD (FDR < 0.1, Figure 5C).

Our analyses showed that circCDYL interacts strongly and specifically with GRWD1 in HepG2. RIP experiments showed that circCDYL also interacts with IGF2BP1 and IGF2BP2. If circCDYL regulates mRNA abundance through its interaction with RBPs, we would expect an enrichment of RBP binding sites in genes affected by circCDYL KD. Based on eCLIP data, we evaluated the presence of GRWD1 and IGF2BP1 binding sites in all genes and found that GRWD1 binding sites are enriched in downregulated genes (P = 8e-14, Chi-square test), while IGF2BP1 binding sites are enriched in upregulated genes (P = 5e-12, Chi-square test, Figure 5D (HepG2)).

GRWD1 is a multifunctional protein (reviewed in (47)), which is overexpressed in cancer cells (48). Specifically, elevated GRWD1 expression has been shown to negatively regulate *TP53* and promote tumorigenesis (49). Since circCDYL is highly covered with GRWD1 binding sites, it could potentially function as a sponge for GRWD1 and lower its binding to targets, incl. *TP53*. We assessed whether circCDYL and GRWD1 regulate the same genes by evaluating the overlap of genes affected by KD. We obtained mRNA expression upon GRWD1 KD in HepG2 from ENCODE (33) and found a significant overlap between altered genes upon circCDYL KD and GRWD1 KD (P < 2.2e-16, Fisher’s Exact Test, Figure 5E (HepG2)). Consistent with an RBP sponge hypothesis, we observed that most downregulated genes upon circCDYL KD are upregulated upon GRWD1 KD (Figure 5F (HepG2)). Accordingly, we found that *TP53* levels are downregulated upon circCDYL depletion in HepG2 cells (P = 0.07, Wald Test, Supp. Figure 5C), while *TP53* expression is slightly upregulated upon GRWD1 KD (P = 0.2, Walt Test, Supp. Figure 5D).

IGF2BP1 is a post-transcriptional regulator that affects the stability, translatability, and localization of essential mRNAs involved in tumor cell proliferation, growth, and invasion (reviewed in (50)). IGF2BP1 is generally assigned oncogenic roles and has been shown to interact with and regulate the expression of genes involved in the MAPK signal transduction pathway (51–53), providing a link to the activation of this pathway upon circCDYL KD. The gene *PLA2G2A* is an extreme outlier upon circCDYL KD with abundance increasing more than 19 times (*P* < 3e-218, Wald Test, Figure 5G). *PLA2G2A* is part of the MAPK pathway, is upregulated in colorectal cancer (54), and is associated with poor clinical outcomes (55, 56). Contrary, *PLA2G2A* is downregulated upon IGF2BP1 KD (P < 3e-53, Wald Test, Supp. Figure 5E), consistent with an oncogenic role of IGF2BP1 and MAPK pathway activation.

Our results suggest that circCDYL interacts with GRWD1 in HepG2 to regulate GRWD1 target genes overall consistent with an RBP sponge hypothesis. Besides, circCDYL could be involved in other regulatory mechanisms to regulate classes of genes and individual transcripts in cancer-related pathways.

### CircCDYL interacts with RBPs in bladder cancer and abundance is associated with overall survival

In a previous study, we found that circCDYL is highly expressed in patients with non-muscle invasive bladder cancer (NMIBC) and correlates positively with good prognosis independently of the parent gene (45). Based on our findings here, we hypothesized that circCDYL possess regulatory functions in bladder cancer (BC) by binding RBPs.

First, to investigate circCDYL-RBP interactions in BC, we performed RIP experiments using the three RBPs confirmed to interact with circCDYL in HepG2 and/or K562: IGF2BP1, IGF2BP2, and GRWD1, in BC FL3 cells. The IPs were validated with western blotting (Supp. Figure 6A) and pull-down of circCDYL (Supp. Figure 6B) and the *CDYL* mRNA (Supp. Figure 6C) were quantified by qRT-PCR (Supp. Table 10). The RIP experiments showed that circCDYL interacts with all three RBPs in BC with higher binding affinity than the host gene (Figure 6A). Contrasting our findings in HepG2, circCDYL pull-down was highly enriched (10x) for IGF2BP1 and IGF2BP2, indicating strong and specific circCDYL-IGF2BPs interactions in BC cells.

**Figure 6:**
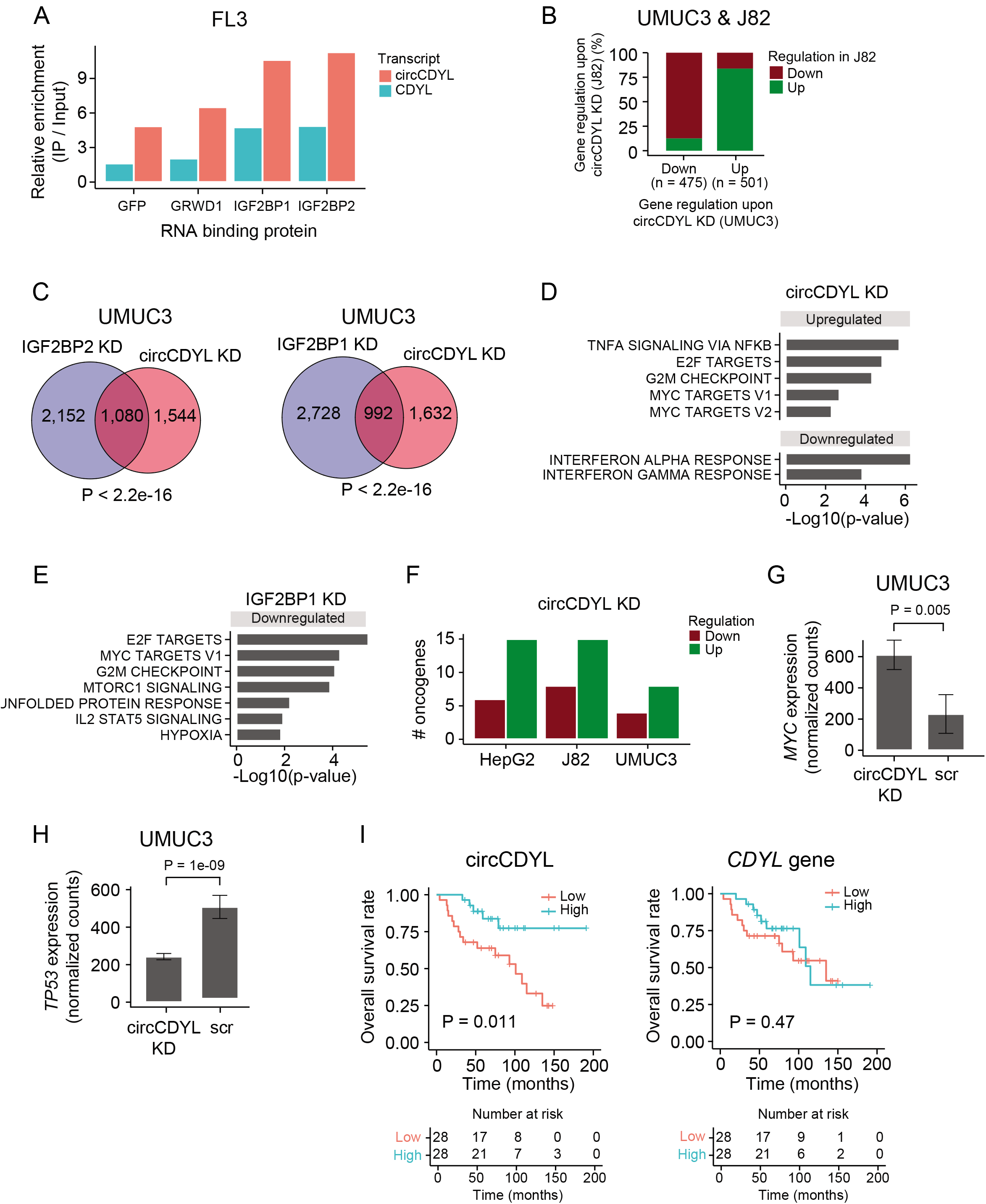
CircCDYL interacts with RBPs in bladder cancer and abundance is associated with overall survival 6A) Relative enrichment of circCDYL and *CDYL* host gene expression levels between immunoprecipitation (IP) and input in the bladder cancer cell line FL3. GFP was used as control. **6B)** Regulation of genes that are differentially expressed upon circCDYL KD in both J82 and UMUC3 cells. 86% of the genes show the same direction of perturbation in both cell lines. **6C)** Overlap of genes affected by circCDYL KD, and IGF2BP1 and IGF2BP2, respectively, in UMUC3. P-values obtained by Fisher’s Exact Test. **6D+E)** Gene set enrichment analysis of 50 hallmarks of cancer upon circCDYL KD (D) and IGF2BP1 KD (E) in UMUC3 cells. **6F)** Distribution of oncogenes among up-and downregulated genes upon circCDYL KD in HepG2, J82, and UMUC3 cells. **6G+H)** Expression of *MYC* (G) and *TP53* (H) upon circCDYL KD in UMUC3. P-values obtained by Wald Test. **6I)** Kaplan-Meier overall survival plots for circCDYL and the host gene in patients with non-muscle invasive bladder cancer. Median expression used as cutoff; circCDYL = 0.125 CPM and *CDYL* = 21.4 FPKM. P-values obtained by Log-Rank Test.

Next, to evaluate the functional role of circCDYL-RBP interactions in BC, we conducted siRNA-mediated KD of circCDYL or the three RBPs (Supp. Table 9). To choose which BC cell lines to use in the experiments, we evaluated the expression of circCDYL using CIRI2 (34) in total RNA-Seq data from eleven BC cell lines (45, 57), and chose J82 and UMUC3 based on cell line stability and high circCDYL expression levels (Supp. Figure 6D). Efficient RBP and circCDYL KD were validated with western blotting (Supp. Figure 6E) and qRT-PCR (Supp. Figure 6F).

From differential expression analysis, we identified 2,624 and 3,014 DE genes upon circCDYL KD in UMUC3 and J82, respectively (Supp. Figure 6G), with a significant overlap of 976 genes (P < 2.2e-16, Fisher’s Exact Test, Supp. Figure 6H). Of these, 86% show the same direction of deregulation in both cell lines, supporting circCDYL-specific alterations across BC cell lines (Figure 6B). When we looked at the genes affected by GRWD1 KD, we found a significant overlap between UMUC3 and J82 (P < 2.2e-16) but not between the BC cells and HepG2 (P > 0.4), consistent with cell-type-specific interactions and functions of circRNAs and RBPs (Supp. Figure 6I). In both BC cells, we found a significant overlap of perturbed genes upon circCDYL KD and depletion of IGF2BP1 and IGF2BP2, respectively (P < 2.2e-16 for all comparisons, Figure 6C, Supp. Figure 6J).

CircCDYL has been suggested to suppress cell growth by inhibiting MYC protein expression levels in bladder cancer through an unknown mechanism (58). IGF2BP1 and IGF2BP2 possess oncogenic roles and positively regulate the expression of various oncogenes (reviewed in (50, 59)). We evaluated how circCDYL and RBP depletion affect 50 hallmarks of cancer (60). We found that several proliferation pathways, e.g. E2F targets, G2M checkpoint, and MYC targets, are upregulated upon circCDYL KD in UMUC3 cells, while genes affecting immune processes are downregulated (FDR < 0.1, Figure 6D). Consistent with a role for circCDYL as a sponge for IGF2BP1 and IGF2BP2, proliferation pathways are downregulated upon IGF2BP1 KD in both UMUC3 (FDR < 0.1, Figure 6E) and J82 cells (FDR < 0.1, Supp. Figure 6K), and immune pathways are upregulated upon IGF2BP2 KD (FDR < 0.1, Supp. Figure 6K). We further examined the distribution of oncogenes (61) among differentially expressed genes upon circCDYL KD, and found twice as many oncogenes among upregulated genes than among downregulated genes in all cell lines (Figure 6F). Accordingly, we found that *MYC* expression is upregulated upon circCDYL KD in UMUC3 cells (P < 0.005, Wald test, Figure 6G) and J82 cells (P = 0.2, Wald test, Supp. Figure 6L). Consistent with our findings in HepG2 cells, we observed that *TP53* levels are downregulated upon circCDYL depletion in both UMUC3 and J82 cells (P < 8e-04, Wald Test, Figure 6H, Supp. Figure 6M).

Finally, we analyzed the expression and clinical correlations of circCDYL in an additional, local cohort of patients with NMIBC (n = 56). Corresponding with our previous findings, high expression levels of circCDYL is positively correlated with overall survival (P < 0.05, Log-Rank Test, Figure 6I) and recurrence-free survival (P < 0.05, Log-Rank Test, Supp. Figure 6N). Expression levels of the *CDYL* host gene is not associated with clinical outcomes (P > 0.4, Log-Rank Test, Figure 6I, Supp. Figure 6N). Additionally, clinical correlations were observed for other circRNAs highly covered with RBP binding sites, e.g. circZKSCAN1 and circRBM33 (P < 0.05, Log-Rank Test, Supp. Figure 6O) and for overall circRNA expression (P = 0.026, Log-Rank Test, Supp. Figure 6P), independent of gene expressions.

Our results show that circCDYL interacts with the oncogenic RBPs IGF2BP1 and IGF2BP2 in BC cells, and that there is a large overlap of altered genes upon circCDYL and RBP depletions. Consistent with a suggested tumor suppressive role, circCDYL depletion activates proliferation processes and the expression of oncogenes. In line with this, we find that elevated expression of circCDYL and other circRNAs identified here to interact with RBPs, are associated with good clinical outcomes.

## Discussion

By comprehensive analyses of a large atlas of eCLIP RBP binding sites and circRNA expression in the ENCODE cell lines HepG2 and K562, we showed that KHSRP binding sites are enriched in introns flanking circRNAs and that KHSRP depletion affects circRNA expression. Additionally, we found that exons comprising circRNAs generally contain more RBP binding sites than non-circularizing exons and that some RBPs preferentially bind to circRNAs. Furthermore, we examined the potential of individual circRNAs to function as RBP sponges and showed experimentally that circRNAs interact with RBPs in a cell-type-specific manner. We specifically investigated the function of circCDYL, which is highly covered with RBP binding sites in both cell lines. We found that circCDYL interacts with GRWD1 in HepG2 cells, and that circCDYL depletion has the opposite effect of knocking down GRWD1. Finally, we showed that circCDYL, which is positively correlated with good clinical outcomes in BC, interacts with IGF2BP1 and IGF2BP2 in BC cell lines, and that circCDYL and RBP KD perturb hallmarks of cancer gene sets, and specifically, that circCDYL KD affects the expression of key tumor genes, e.g. *TP53* and *MYC*.

First, we evaluated the overall potential of circRNAs as a group to interact with RBPs. No previous studies have comprehensively catalogued circRNA-RBP interactions using experimental data. In a previous study, You et al. found that circRNAs are not more prone to bind RBPs than linear mRNAs based on computational nucleotide sequence prediction of RBP binding sites (30). However, due to contextual RBP binding preferences and the unique structures of circRNAs, nucleotide sequences are likely not adequate to predict circRNA-RBP interactions. Here, we defined RBP binding sites transcriptome-wide using experimental eCLIP data and found that RBP-coverage generally increases with transcript abundance. To ensure that enrichment of RBP binding sites in circRNAs is not simply explained by transcript abundance, we compared the RBP-coverage of circRNAs to comparable gene sets. Contrasting the findings by You et al., our analyses showed that circRNAs are highly enriched with RBP binding sites compared to linear exons in host genes and genes of the same expression. Even compared to linear exons in the most highly expressed genes, circRNAs were enriched with or covered by an equal amount of RBP binding sites. These observations indicate specific circRNA-RBP interactions independent of transcript abundance and that non-or inefficiently translated circRNAs are ideal binding platforms for RBPs, since they are not in direct competition with elongating ribosomes.

Since the eCLIP data set was generated for linear transcripts, we evaluated the ability of circRNAs to interact with RBPs across the BSJ by mapping unmapped eCLIP reads to a reference set of BSJ sequences. We identified potential backsplicing events covered by eCLIP reads fulfilling our cut-off (Methods), of which a few conform to circRNAs identified in circBase. Nevertheless, none of them correspond to the BSJ of a circRNA expressed in HepG2 or K562. The lack of eCLIP reads spanning circRNA BSJs could be explained by short eCLIP sequencing reads around 20-25 bp long. This is considerably shorter than reads from most total RNA-Seq data sets, which typically use paired-end libraries and obtain longer read lengths (>= 100 bp). Hence, these are not ideal for robust mapping across the BSJs of circRNAs (62). Apparent backsplicing sequences supported by eCLIP reads could be artefacts or stem from other mechanisms such as template switching by reverse transcriptase, tandem duplication, structural variation between individuals, and RNA trans-splicing (63). They could potentially also be real circRNAs not detected by CIRI2.

RBP binding sites in circRNA loci could reflect RBPs binding to the circRNA, the corresponding linear transcript, or both. Additionally, many factors could influence RBP binding capacity, like expression of cofactors and secondary structures of circRNAs. To confirm the validity of predicting circRNA-RBP interactions from eCLIP data, we experimentally validated a set of circRNA-RBP interactions by RIP. All predicted circRNA-RBP interactions were validated except for one. Negative results would be expected if the RBP only interacts with the linear transcript and not the circRNA or if the circRNA-RBP interaction only occurs under certain circumstances.

By analyzing the RBP-coverage of abundant circRNAs, we found that circCDYL is almost completely covered with RBP binding sites in both cell lines. In HepG2, GRWD1 binding sites cover 93% of the exonic circCDYL sequence, which is comparable to the efficient miRNA sponge, ciRS-7, which is ~90% covered by motifs comprising miR-7 binding sites (64). RIP experiments showed that circCDYL interacts with GRWD1, IGF2BP1, and IGF2BP2 in both HepG2 and BC cells. From eCLIP data, we observed no IGF2BP1 binding sites in circCDYL in HepG2, however, we found that circCDYL was pulled down more efficiently by both of the IGF2BPs than by GRWD1 in BC cells. Different binding affinities, competition for binding sites, dynamic interactions, and expression of RBPs and cofactors could influence interactions and explain the different observations in HepG2 and BC cells.

CircCDYL is clinically interesting, because it is deregulated in diseases (9, 44, 45), correlated to clinical outcomes (45), and highly expressed in patient plasma samples (44) and exosomes (9), indicating potential as a non-invasive biomarker. Sun et al. suggested that circCDYL overexpression inhibits MYC at the protein level, but found no effect on *MYC* mRNA levels (58), contrasting our findings here. Since circCDYL is almost completely covered with RBP binding sites, we speculate that circCDYL regulates important tumor genes and hallmarks of cancer through RBP interactions. Further experiments should address the regulatory potency of circCDYL on tumorigenesis and assess its clinical value as a new biomarker. Despite clinical correlations, circRNA perturbation in cancer might not be causative but a consequence of underlying biological mechanisms.

The interplay between circRNAs and RBPs are highly complex as both factors have been shown to modulate the function and expression of the other. Zhang et al. implied that circRNA localization might be facilitated by RBP-mediated transportation in HepG2 (39). Many of the RBPs we identified with enriched binding sites in circRNAs occupy both the nuclear and cytoplasmic fraction of the cell (Supp. Table 5) and could be involved in circRNA nuclear export or otherwise in circRNA localization. For instance, the IGF2BPs, which we found on average have more target sites in circularizing exons, are regulators of essential mRNAs in tumorigenesis and are important players of mRNA stability and transportation to subcellular compartments (59). Accordingly, IGF2BP3 was found to bind to several circRNAs (29). Additionally, RBPs can regulate circRNA biogenesis by binding to intronic regions under certain circumstances. Here, we found that KHSRP depletion affects the expression of a subset of circRNAs with KHSRP binding sites in flanking intronic regions. Finally, RBPs binding to circRNAs could mark the circRNAs as “self” to prevent innate immune responses as suggested by Chen et al. (65).

CircRNAs can also regulate RBP localization and function. Binding of specific RBPs across multiple circRNAs and overall clinical correlations might reflect large-scale regulatory roles of circRNAs as a group. Additionally, circRNAs could act as dynamic scaffolds that bring together regulatory complexes. CircRNAs enriched with RBP binding sites, e.g. circCDYL, could bind and sequester a large number of RBPs to certain subcellular compartments and abrogate their normal function similarly to the role of the miR-7 sponge, ciRS-7 (24, 64). Our results showed that circRNAs interact with RBPs in a highly cell-type-specific manner, consistent with findings for circFoxo3, which bind diverse RBPs in different biological settings (27, 28, 66). CircRNAs might obtain different tertiary structures in various tissues and cellular conditions (67), partly explaining diverse functions and dynamic circRNA-RBP interactions as observed for circCDYL.

The list of potential functions of circRNA-RBP interactions is long and their regulatory dependencies complicated. How circRNAs mechanistically controls RBP localization, activity, and homeostasis awaits further investigation, but here we have provided strong comprehensive and global-scale evidence to support such roles.

## Material and Methods

### HepG2 and K562 encode cell lines

We downloaded all total and fractionated samples for the cell lines HepG2 (n = 5) and K562 (n = 11) from ENCODE (https://www.encodeproject.org/), generated by the labs of Brenton Graveley, UConn; Thomas Gingeras, CSHL; and Éric Lécuyer, IRCM (Supplementary file 1+2).

### Detection of circRNAs in HepG2 and K562

We used the CIRI2 pipeline (v2.0.6) (34) to detect circRNAs in all samples. Before running the CIRI2 pipeline we trimmed reads with Trim Galore and cutadapt (v0.4.1 and v1.9). We aligned the reads to the human genome (hg19) using bwa (v0.7.15) and samtools (v1.3). The CIRI2 pipeline was run with a gene transfer format (GTF) file (hg19) to annotate the overlapping gene of the circRNAs. The pipeline does not take strand into consideration so afterwards we named the overlapping gene of the circRNAs according to strand information.

### Gene expression profiling in HepG2 and K562

Illumina paired end (2x 50bp reads) were stripped of library adapters (Trim Galore and cutadapt as above) and mapped to the human genome (hg19) using TopHat2 (version 2.1.1) (68) and Bowtie2 (version 2.2.8.0) (69). Cufflinks (v2.1.1) (70) and HTSeq (v0.6.1p1) (71) were used to estimate the transcripts abundance using transcript information from GENCODE v19. Samtools (v1.3) (72) and Picard (v2.0.1) were used for quality control and statistics.

### eCLIP of HepG2 and K562

The eCLIP data of RNA binding protein (RBP) targets from HepG2 and K562 was obtained from a previous study (32). Briefly, RNA-RBP interactions were covalently linked by UV, followed by RNA fragmentation and immunoprecipitation (IP) with a specific antibody against the RBP of interest. RNA from RNA-RBP complexes were prepared into paired-end high-throughput sequencing libraries and sequenced. All experiments were performed with a size-matched input control (SMInput), which control for nonspecific background signal. RBP targets were determined using the CLIPper algorithm (73). To report RBP target sites, IP fold enrichment were calculated based on the number of reads overlapping peaks identified by CLIPper in both the IP and SMInput samples. A cut-off of log2(foldchange) >= 3 was applied, which means an 8-fold enrichment in the IP. Additionally, we only considered RBP binding sites supported by at least 10 reads in the IP.

### eCLIP reads spanning BSJ

We generated a reference set of all possible exonic BSJ events within transcripts by extracting 30 bp from each side of all exons and pasting them together in reverse order. Unmapped eCLIP reads (~20-25 bp) from both IP and input experiments (32) were mapped against the BSJ reference set, allowing two mismatches. We removed PCR duplicates based on the barcode and position of reads and merged barcode replicates. Before counting mapped reads spanning backsplice junctions, we removed read 1 as this was only used to identify PCR duplicates. Only reads spanning the BSJ site by >= 5 bp were considered. To identify specific RBP binding sites, we used the same cut-off as above: only RBP binding sites supported by >= 10 reads in both IP replicates and with a log2(foldchange) >= 3 between IP and input were reported. A pseudo count of 1 was added before counting the foldchange.

### Subcellular localization of RBPs in HepG2

Localization of RBPs by immunofluorescence was performed for another study (38). We only considered cellular fractions where we had both circRNA and RBP data, e.g. the nuclear and cytoplasmic fractions of HepG2.

### Genomic annotations of exons

We extracted all exons from known and novel protein coding transcripts (hg19). Exons from different transcript isoforms were merged using bedtools merge (v2.25.0) to obtain one genomic interval per exon per gene. We used bedtools intersect to obtain exonic regions of the circRNAs. Exons in circRNAs were annotated as backsplice junction (BSJ) circ-exons if they are involved in the backsplicing event and circ-exons if they are internal exons in the circRNAs. An exon might be an internal circ-exon in one circRNA but a BSJ circ-exon in another circRNA. In that case, it will be annotated as a BSJ circ-exon. Intronic regions were achieved in the same way.

### RBP binding sites in circRNAs

We used bedtools intersect to annotate the overlap between the exonic parts of circRNAs and RBP binding sites. We disregarded RBPs with a binding site < 4 bp or a circRNA-RBP overlap < 4 bp long. To get that fraction of circRNAs covered with RBP binding sites, we merged the RBP binding sites overlapping circRNAs and intersected them with merged exons in circRNAs.

### RBP binding sites transcript-wide

We extracted all merged exon positions (of known and novel protein coding transcripts from above) for expressed genes in each cell line and intersected them with RBP binding sites. Again, we disregarded RBPs with a binding site < 4 bp or an exon-RBP overlap < 4 bp. We obtained the fraction of exons covered with RBP binding sites by merging these RBP binding sites and intersecting them with merged exon positions from expressed genes.

To compare RBP-coverage of circRNAs to comparable genes sets, the genes were divided into 100 bins based on expression and gene expression were considered a proxy for exon expression. We randomly drew internal non-circ-exons from genes not producing circRNAs, matching the gene expression distribution of genes producing highly expressed circRNAs, with one hundred repetitions. The same approach was conducted to extract exons of highly expressed genes (percentile 90-95 and 96-100).

To evaluate the enrichment of circRNA RBP-coverage to non-circ-exons in host genes, we added a pseudo count of 1 to the RBP-coverage of circRNAs and non-circ-exons to include circRNAs and non-circ-exons with no RBP binding sites in the analysis. We divided circRNA RBP-coverage by the mean RBP-coverage of internal non-circ-exons within host genes. CircRNAs from genes with zero internal non-circ-exons are not depicted in the figure and are denoted with NA in Supp. Table 7.

### RBP binding sites in introns flanking circRNAs and non-circularizing exons

RBP binding sites in intronic regions were evaluated in the same way as exonic RBP binding sites (above). Intronic regions overlapping exonic regions in a transcript isoform were omitted. To identify RBPs with binding sites in both flanking introns of circRNAs, flanking regions (10,000 bp to each side of the circRNAs) were intersected with RBP binding sites in intronic regions. For non-circularizing exons, the first and last exons from expressed genes were removed from this analysis as they are not flanked by introns on both sides. Additionally, exons found in circRNAs were disregarded. As for the circRNAs, flanking regions of internal non-circularizing exons (10,000 bp to each side) were intersected with RBP binding sites in intronic regions.

### Alu repeats in introns

Positions of *Alu* repeats were obtained from the UCSC Browser RepeatMasker track as described in a previous study (45). *Alu* repeat positions were intersected with flanking regions of circRNAs and non-circularizing exons from above.

### Gene Ontology enrichment analysis

Gene Ontology (GO) term data was extracted from BioMart (74). Two subsets of the eCLIP data were generated: RBPs that bind relatively more to BSJ circ-exons than non-circ-exons and a negative set containing all other RBPs for each of the two cell lines. In K562, the subset of RBPs that are more prone to bind to BSJ circ-exons (n = 12) contains NOLC1, IGF2BP1, IGF2BP2, YBX3, RBM15, UCHL5, LIN28B, PUM1, ZNF800, GRWD1, U2AF2, and ZNF622. In HepG2, the subset (n = 8) contains BAP1, BCLAF1, ZNF800, GRWD1, RBM15, SF3A3, TRA2A, and IGF2BP3. Only GO terms represented by at least two RBPs were considered in the analysis for each cell line. For the GO terms in each of the GO domains; Biological process (BP), Cellular component (CC) and Molecular function (MF), a one-tailed Fisher’s exact test was performed.

### Pathway analyses and distribution of oncogenes

We used the R package *gage* (75) for gene set enrichment and pathway analyses. We used the gene sets provided by the package for KEGG pathway analyses. We obtained the 50 hallmarks of cancer gene sets from The Molecular Signatures Database (MSigDB) (60). We downloaded the list of all cancer census genes from The Cosmic Cancer Gene Census (61). Some genes were classified into several categories, so we restricted our analyses to genes that were only classified as oncogenes.

### RIP assay for HepG2 and K562 cells

Five RBPs and four circRNAs were chosen for validation. We designed primers against the unique backsplice junction of specific circRNAs to validate their expression in HepG2 and K562 (Supplementary Table 1). Antibodies were obtained for the following five RBPs: GRDW1 (Bethyl A301-576A Lot 1), UCHL5 (Bethyl A304-099A Lot 1), YBX3 (A303-070A Lot 1), IGF2BP1 (MBL RN007P Lot 004), and IGF2BP2 (MBL RN008P Lot 005). Rabbit IgG Isotype Control (Invitrogen Cat# 02-6102) were used as negative control.

K562 cells (American Type Culture Collection (ATCC^®^), CCL-243^™^) were grown in RPIM 1640 media (Gibco^™^, Life Technologies, 11875119) with 10% FBS (Gibco^™^ Life Technologies, 26140079). HepG2 cells (American Type Culture Collection (ATCC^^®^^), HB-8065) were grown in DMEM (Gibco^™^, Thermo Fisher Scientific) with 10% FBS (Gibco^™^ Life Technologies, 26140079). 20 × 10^6^ snap frozen cells were lysed in 1ml of iCLIP Lysis Buffer (50mM Tris-HCl pH 7.4, 100mM NaCl, 1% NP-40 (Igepal CA630), 0.1% SDS, 0.5% Sodium Deoxycholate) with 5.5μl Protease Inhibitor Cocktail Set III EDTA Free (EMD Millipore Corp.539134-1ML) and 11μl Murine RNase Inhibitor (New England BioLabs Inc^®^ M0314L) for 15 mins and were then centrifuged at 20,000 g for 20 mins at 4°C. The supernatant was placed in a solution containing specific primary(10μg)/secondary(1.25g) (anti-Rabbit magnetic DynaBeads, Invitrogen, 11204) antibody-antibody (incubated on a rotator at 25°C for 45 minutes) to immunoprecipitate overnight on a rotator at 4°C. The RNA-RBP pull down was then purified by stringently washing with NET-2 wash buffer (5mM Tris-HCl pH 7.5, 150mM NaCl, 0.1% Triton X 100). Isolation of RNA from the RNA-RBP complexes were accomplished with the addition of TRIzol^™^ Reagent (Invitrogen^™^, 15596018) followed by the Direct-zol^™^ RNA MiniPrep (Zymo Research Cat No. R2052). Isolated RNA was reverse transcribed with SuperScript^™^ III First Strand Synthesis System (Invitrogen^™^ 18080051) using Random Hexamers (ThermoFisher Scientific N8080127) and circRNAs were amplified using GoTaq^®^ DNA Polymerase (Promega, M3005); 4μl of 1:2.5 diluted cDNA, 1μl of each primer for 34 cycles at the following conditions: strand separation at 95°C for 30s, primer hybridization at 55°C for 30s, and elongation at 72°C for 20s followed by a final elongation step at 72°C for 5 min. Amplicons were run on a 3% Agarose Gel at 135V for 35 minutes at 4°C alongside a 50bp ladder marker. CircRNAs bound by RBPs were identified on the gel based on amplicon size (Supplementary Table 1).

### Validation of RBP Pulldown in IP

Approximately 5 × 10^6^ cells from the overnight incubation with the primary/secondary antibody-antibody complex was stringently washed with NET-2 buffer then denatured in a DL-Dithiothreitol (DTT) (Sigma-Aldrich D9779), NuPAGE^™^ LDS Sample Buffer (4X) (Invitrogen^™^ NP0008) mixture at 70°C for 10 min in a Thermomixer at 1200 rpm. Denatured protein samples were subject to SDS-PAGE and western blotting with Anti-Rabbit IgG HRP (Rockland Inc. 18-8816-33).

### RBP knockdown in HepG2

RBP knockdown samples were originally generated for the ENCODE project by Brenton Graveley’s Lab, UConn (33). RNA-Seq data of polyadenylated transcripts were obtained from ENCODE (https://www.encodeproject.org/).

### siRNA-mediated knockdown

3×10^5^ cells (UMUC3, J82 or HepG2) were seeded per well in 6-well plates (2 ml per well). 24 hrs later cells were transfected with siRNA (Supp. Table 9) using SilentFect (Bio-Rad) in biological triplicates: 100 μl OptiMEM (Gibco) + 2 μl SilentFect (Bio-Rad) was incubated at RT for 5 min before mixing with 2 μl siRNA (20 μM stock) pre-diluted in 100 μl OptiMEM. After gentle mixing by pipetting, the solution was incubated for 20 mins at RT prior to dropwise addition to cells (final concentration of 20 nM). Cells were incubated for 56 hrs and harvested by addition of 1 ml Trizol (Thermo Fisher Scientific) to each well followed by mixing. RNA was purified according to manufacturer’s protocol with an additional chloroform extraction to increase quality. A parallel set of identically treated samples were used to harvest cells for protein lysates by addition of 300 μl 2 X SDS sample buffer [4% SDS, 20% glycerol, 10% 2-mercaptoethanol, 0.004% bromophenol blue and 0.125 M Tris-HCl, pH 6.8] per well followed by incubation at 90°C for 8-10 min until all cell material is dissolved. Knockdown efficiency was assessed either by Western blotting or qRT-PCR.

For qRT-PCR, RNA samples were treated with DNase I (Thermo Fisher Scientific) according to manufacturer’s protocol. First-strand cDNA synthesis was carried out using the Maxima First Strand cDNA synthesis Kit for qPCR (Thermo Fisher Scientific) according to manufacturer’s protocol. qPCR reactions were prepared using gene-specific primers and Platinum SYBR Green qPCR Supermix-UDG (Thermo Fisher Scientific) according to manufacturer’s protocol. An AriaMx Real-time PCR System (Agilent Technologies) was used for quantification of RNA levels and the X_0_ method was used for calculations of relative RNA levels (76) normalized to GAPDH or beta-actin mRNA as indicated.

For western blotting, Cell lysates dissolved in SDS load buffer were heated at 90°C for 3 min and separated on a Novex WedgeWell 4-12% Tris-Glycine Gel (Invitrogen). Proteins were transferred to a PVDF Transfer Membrane (Thermo Scientific) using standard procedures. The membranes were blocked in 5% skimmed milk powder in PBS for 1 hour at room temperature. The membranes were incubated at 4°C overnight with primary antibodies diluted as indicated in 5% skimmed milk powder in PBS. After three times wash in 13 ml PBS, the membranes were incubated with goat polyclonal HRP-conjugated secondary antibodies (Dako) diluted 1:20000 in 5% skimmed milk powder in PBS. After 1 hour of incubation at room temperature, the membranes were washed three times in 13 ml PBS and the bound antibodies were detected using the SuperSignal West Femto maximum sensitivity substrate (Thermo Scientific) according to manufacturer’s protocol and the LI-COR Odyssey system (LI-COR Biosciences).

We quantified mRNA expression using QuantSeq (46). Sequencing libraries were generated using the QuantSeq 3’mRNA Library Prep Kit Protocol (Lexogen). Input were 500 ng RNA and 12 PCR cycles were applied. Library concentrations were measured on Qubit 3.0 (invitrogen) and the average size of the library was measured on TapeStation. The libraries were 1×75 bp sequenced on a NextSeq500 system (Illumina). QPCR Universal Human Total Reference RNA (UHR) (Agilent, Cat no: 750500) was included in all batches to assess batch effect.

The raw reads were converted to fastq format and demultiplexed using Illumina’s bcl2fastq v2.20.0.422 and library adapters were removed from the read pairs (trim_galore v0.4.1). Reads were mapped to the human genome (hg19) using TopHat2 (version 2.1.1) (68) and Bowtie2 (version 2.1.0.0) (69), and Cufflinks (v2.1.1) (70) and HTSeq (v0.11.2) (71) were used to estimate the transcripts abundance using transcript information from GENCODE v19. Samtools (v1.3) (72) and Picard (v2.0.1) were used for quality control and statistics.

### RIP assay for FL3 cells

1.65×10^6^ FL3 cells were seeded in a P10 dish and transfected 24 hrs later with 12 μg total DNA (2 μg Twin-Strep-Tag-RBP expression vector and 10 μg pcDNA3 PL). 48 hrs later cells were washed with PBS and placed on ice. 1 mL cold lysis buffer (50 mM TRIS-HCl pH 7.5, 10 mM NaCl, 5 mM MgCl_2_, 0.5% Triton-X 100, 2% Hexane-1,6-diol, 1 pill Complete pr. 10 mL) was added per plate and cells were scraped off and transferred to an Eppendorf tube. Samples were mixed and spun (13,000 RPM, 15 min at 4ºC) and 50 μL supernatant was transferred to 50 μL SDS load buffer while 100 μL supernatant was transferred to 0.5 mL Trizol (INPUT). 800 μL supernatant was incubated with pre-equilibrated MagStrep type 3 XT beads (IBA Life sciences) and rotated for ≥2 hrs at 4ºC. 500 μL supernatant was collected (FT) and samples were washed 4x with 1.5 mL WASH1 buffer (10 mM TRIS-HCl pH 7.5, 150 mM NaCl, 5 mM MgCl_2_, 0.1% Triton-X 100) and 2x with WASH2 buffer (10 mM TRIS-HCl pH 7.5, 150 mM NaCl, 5 mM MgCl_2_). During the last wash sample beads were divided 1:10 (RNA:protein) and added 40 μL SDS load buffer or 0.5 mL Trizol (IP). INPUT, FT and IP samples were analyzed with western blotting and RT-qPCR.

### Total RNA-Seq of KHSRP KD in HepG2 and K562

Total RNA from KHSRP knockdown and control samples were obtained from Brenton Graveley’s Lab, UConn (33). Total RNA sequencing libraries were prepared using the KAPA RNA HyperPrep Kit with RiboErase for Illumina Platforms (KAPA Biosystems). Briefly, DNA oligos were hybridized to rRNA and digested using RNase H treatment followed by a 2.2x bead-based cleanup. Next, RNAs hybridized to rRNA targeting oligos were removed from the samples via RNase H digestion, followed by a 2.2x bead-based cleanup. The rRNA-depleted RNA was then fragmented to an insert size of 200-300 bp at 94 °C for 6 minutes in the presence of Mg2+. 1st strand cDNA was synthesized using random primers followed by 2nd strand dscDNA synthesis marked by dUTP and A-tailing with dAMP at the 3’-end. 3’-dTMP adapters were ligated to the 3’-dAMP library fragments. A 0.63x bead-based cleanup followed by a 0.7x bead-based cleanup were performed and the purified, adapter-ligated library DNA was amplified with 12 amplification cycles followed by a 1x bead-based cleanup. Post-capture libraries were barcoded and pooled for sequencing. Libraries were sequenced on an Illumina HiSeq 4000 instrument to a depth of approximately 40 million 100 bp paired-end reads. CircRNA expression was profiled using CIRI2 as described for HepG2 and K562 with the following tool versions: Trim Galore (v0.4.1), cutadapt (v1.15), bwa (v0.7.17), and samtools (v1.9).

### Bladder cancer patient cohort

RNA was paired end sequenced using an Illumina NextSeq 500 instrument. Reads were demultiplexed using bcl2fastq v2.18.0.12 trimmed for traces of adapters using Trim Galore v0.4.1, and mapped to the hg19 genome build using tophat v2.1.1. Gene expression was estimated using cufflinks v2.1.1 and HTseq v0.6.1. CircRNA expression was quantified using the CIRI2 pipeline as described above for HepG2 and K562.

### Statistical analyses

All statistical tests were performed in R (77, 78). The non-parametric Wilcoxon Rank Sum Test was utilized to evaluate the enrichment of RBP binding sites in circularizing exons compared to non-circularizing exons from different groups. It was also applied to compare the enrichment of RBP binding sites in circularizing exons to non-circularizing exons for individual RBPs. The chi-square test was assessed to evaluate enrichment of individual RBPs in flanking introns of circRNAs compared to non-circularizing exons. The empirical p-value was calculated to evaluate the significance of a specific observation compared to the distribution of all observations. *DESeq2* (79) was used for differential expression analyses of circCDYL and RBP knockdown data. The R packages *Survival* (80, 81) and *Survminer* (82) were used to produce Kaplan-Meier plots and curves were compared statistically by the log-rank test. For multiple testing corrections, we applied Benjamini-Hochberg correction and statistical differences were declared significant at FDR < 0.1. When multiple testing was not applied, statistical differences were declared significant at P < 0.05. Most of the plots were produced with the R package *ggplot2* (83).

## Declarations

Informed written consent to take part in future research projects was obtained from all patients, and the specific project was approved by the National Committee on Health Research Ethics (#1708266).

### Consent for publication

Not applicable.

### Availability of data and materials

All generated sequencing data have been deposited in NCBI’s Gene Expression Omnibus. The circCDYL knockdown sequencing data in HepG2, UMUC3, and J82 cells and the GRWD1, IGF2BP1, and IGF2BP2 knockdown sequencing data in UMUC3 and J82 cells are accessible through accession number GSE146726, https://www.ncbi.nlm.nih.gov/geo/query/acc.cgi?acc=GSE146726. Total RNA-Seq data upon KHSRP knockdown is accessible through accession number GSE145984, https://www.ncbi.nlm.nih.gov/geo/query/acc.cgi?acc=GSE145984. Sensitive personal data (raw data from RNA-Seq and clinical information) cannot be shared publicly because of Danish legislation regarding sharing and processing of sensitive personal data. The publicly available datasets supporting the conclusions of this article, e.g. eCLIP data, total RNA-Seq data from HepG2 and K562 etc., are available in the ENCODE repository, https://www.encodeproject.org/, as described in the Methods sections.

### Competing interests

The authors declare that they have no competing interests.

### Funding

TLHO and JSP were supported by generous grants from the Lundbeck Foundation (#R191-2015-1515), the Danish Cancer Society (R124-A 7869-15-82), the Danish Council for Independent Research | Medical Sciences (FSS; DFF - 7016-00379), the Harboe Foundation (19110), and Aage and Johanne Louis-Hansen Foundation (19-2B-503). ABK and CKD were supported by the Danish Cancer Society (R167-A11105). We thank the following funds for supporting TLHO’s stays in GY’s lab and for supporting the experimental follow-up studies: the Harboe Foundation (18004), the Danish Cancer Society (R215-A12954), the Family Hede Nielsen’s Foundation, the Lundbeck Foundation (R296-2018-2094), the Graduate School of Health at Aarhus University, and the NEYE Foundation. Clinical sample procurement was supported by the Danish Cancer Biobank, and LD was supported by The Danish Cancer Society (R204-A12270).

### Authors’ contributions

TLHO and JSP conceived the project. TLHO performed the data analyses and generated all figures. SP performed RNA immunoprecipitation in HepG2 and K562 cells, while CKD and ABK performed RNA immunoprecipitation in FL3 cells. CKD and ABK also conducted siRNA-mediated knockdown of circCDYL in HepG2, J82, and UMUC3 cells and of RBPs in J82 and UMUC3 cells. AS prepared total RNA-Seq libraries from KHSRP knockdown samples. SS generated the BSJ reference set and contributed initial analyses of eCLIP data. SV profiled circRNA and gene expression in RNA-Seq data sets. AMR performed GO enrichment analysis. MMN contributed insights into expression analyses. NF and LD generated and contributed clinical cohorts. SA and GY contributed eCLIP data and supervised analyses. JSP supervised the project. TLHO and JSP wrote the manuscript with input from the other authors.

## Acknowledgements

We thank Brenton Graveley for providing total RNA from KHSRP knockdown samples.

## Supplementary figure legends

**Supplementary Figure 1:**
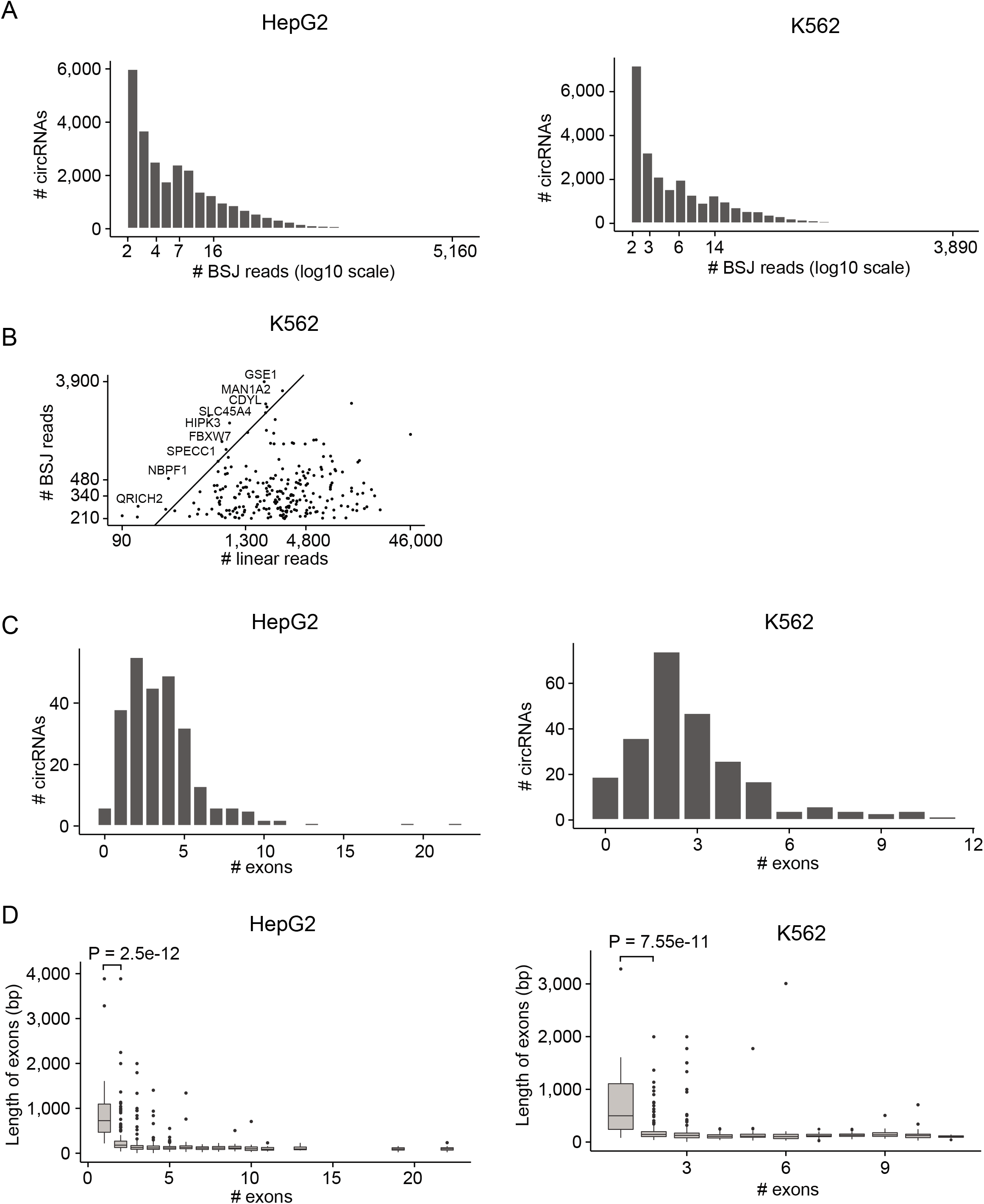
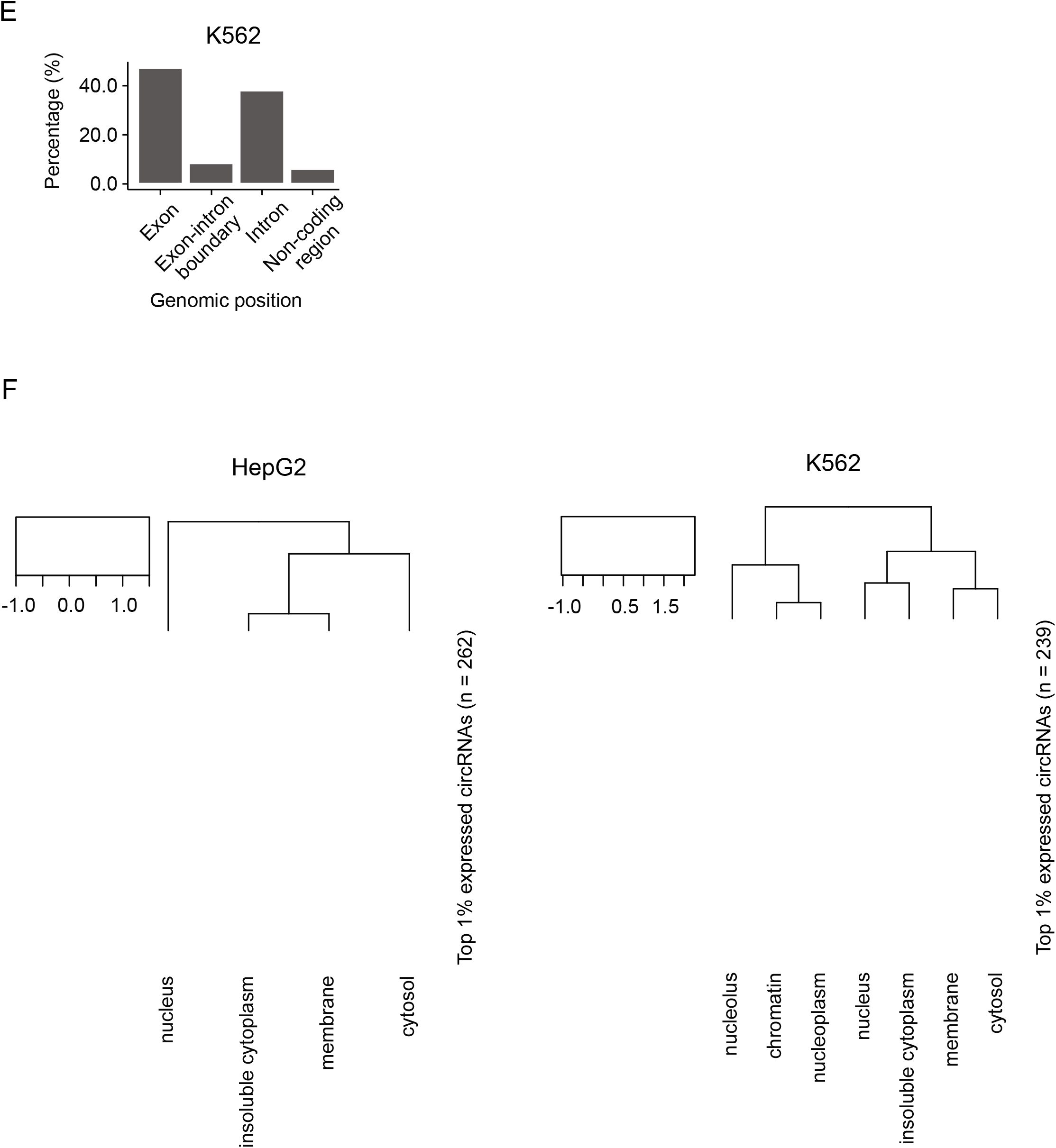
Circular RNAs are highly expressed in HepG2 and K562 and generally colocalizes with RBPs. **1A)** Number of backsplice junction (BSJ) spanning reads supporting circRNAs in HepG2 (left, n = 26,174) and K562 (right, n = 24,076). X-axis is plotted on a logarithmic scale (log10). **1B)** Number of backsplice junction (BSJ) reads supporting the 1% highest expressed circRNAs (n = 241) and the number of reads spanning canonical splice sites in the corresponding linear transcripts in K562. X-and Y-axis are plotted on a logarithmic scale (log10) showing actual counts. Some circRNAs are depicted with their host gene name. **1C)** Number of exons comprising highly expressed circRNAs in HepG2 (left) and K562 (right). 74% (HepG2) and 83 % (K562) of the top 1% expressed circRNAs contain less than 5 exons, while 14% (HepG2) and 15% (K562) of the circRNAs originate from just one exon. The x-axis shows the number of exons within the circRNA loci. **1D)** Length of exons comprising circRNAs in HepG2 (left) and K562 (right). The x-axis shows the number of exons within the circRNA loci. **1E)** Genomic location of RBP binding sites in circRNA loci in K562. Most RBP binding sites are found within exonic parts of the circRNAs. **1F)** Heatmap of circRNA expression in subcellular fractions of HepG2 (left, n = 4) and K562 (right, n = 7). Two of the top 1% highest expressed circRNAs in K562 were not identified in the subcellular fractions.

**Supplementary Figure 2:**
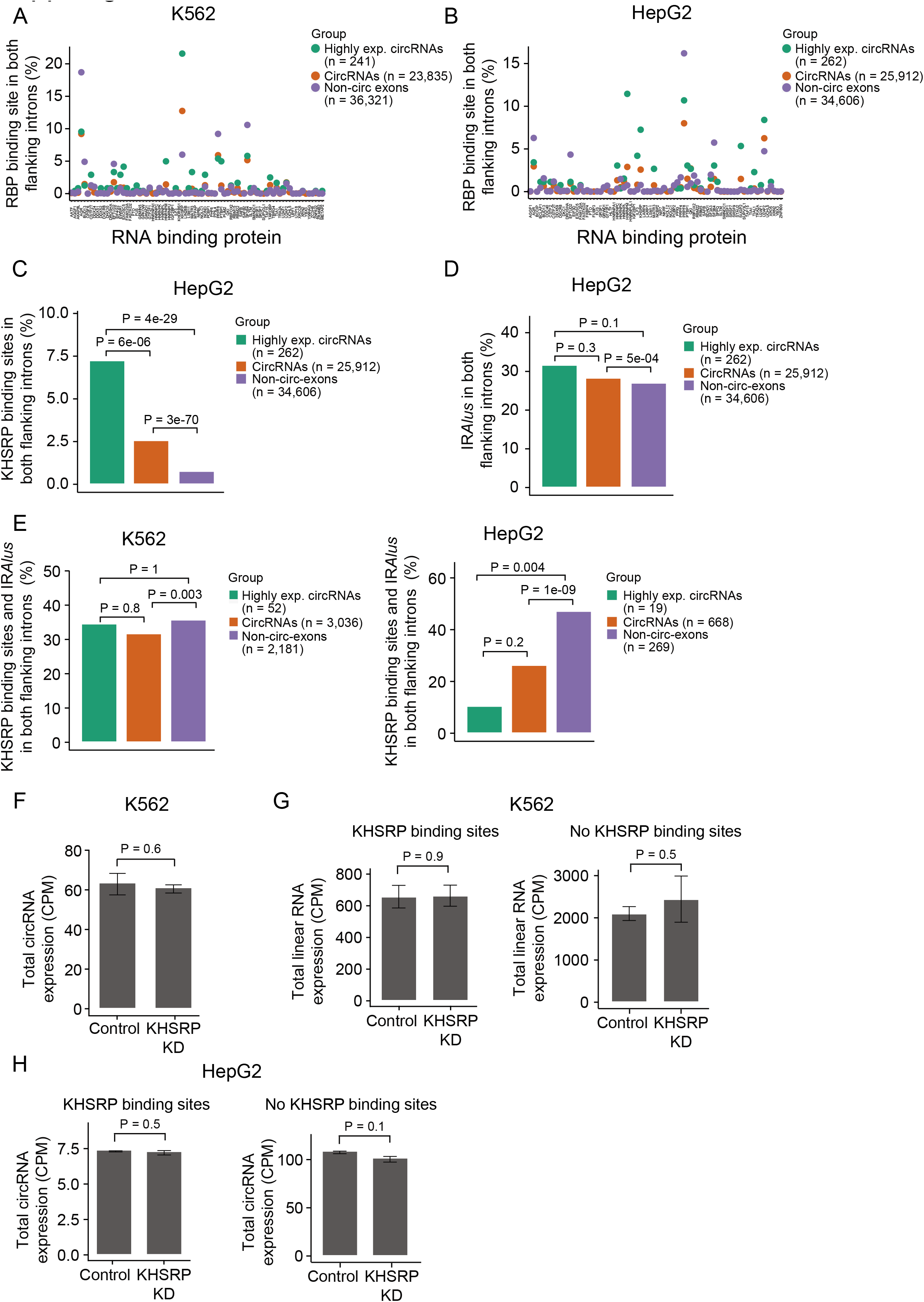
KHSRP binding is enriched in introns flanking circRNAs and affects biogenesis 2A+B) Percentage of circRNAs and non-circ-exons with RBP binding sites in both flanking introns for each RBP in K562 (A) and HepG2 (B)**. 2C)** Percentage of circRNAs and non-circ-exons in genes producing circRNAs with KHSRP binding sites in both flanking introns in HepG2. P-values obtained by Chi-square Test. **2D)** Percentage of circRNAs and non-circ-exons in genes producing circRNAs with inverted repeated *Alu* elements (IR *Alus*) in both flanking introns in HepG2. P-values obtained by Chi-square Test. **2E)** Percentage of circRNAs and non-circ-exons in genes producing circRNAs with KHSRP binding sites in both flanking introns that are also surrounded by IR *Alus* in K562 (left) and HepG2 (right). P-values obtained by Chi-square Test. **2F)** Total circRNA expression (n = 1,834) in KHSRP knockdown (KD) and control samples in K562. P-values obtained by T-test. CPM = Counts per million. **2G)** Total expression of the corresponding linear RNAs of circRNAs with (left, n = 297) and without (right, n = 1,537) KHSRP binding sites in both flanking introns in KHSRP knockdown (KD) and control samples in K562. P-values obtained by T-test. **2H)** Total expression of circRNAs with (left, n = 97) and without (right, n = 2,541) KHSRP binding sites in both flanking introns in KHSRP knockdown (KD) and control samples in HepG2. P-values obtained by T-test.

**Supplementary Figure 3:**
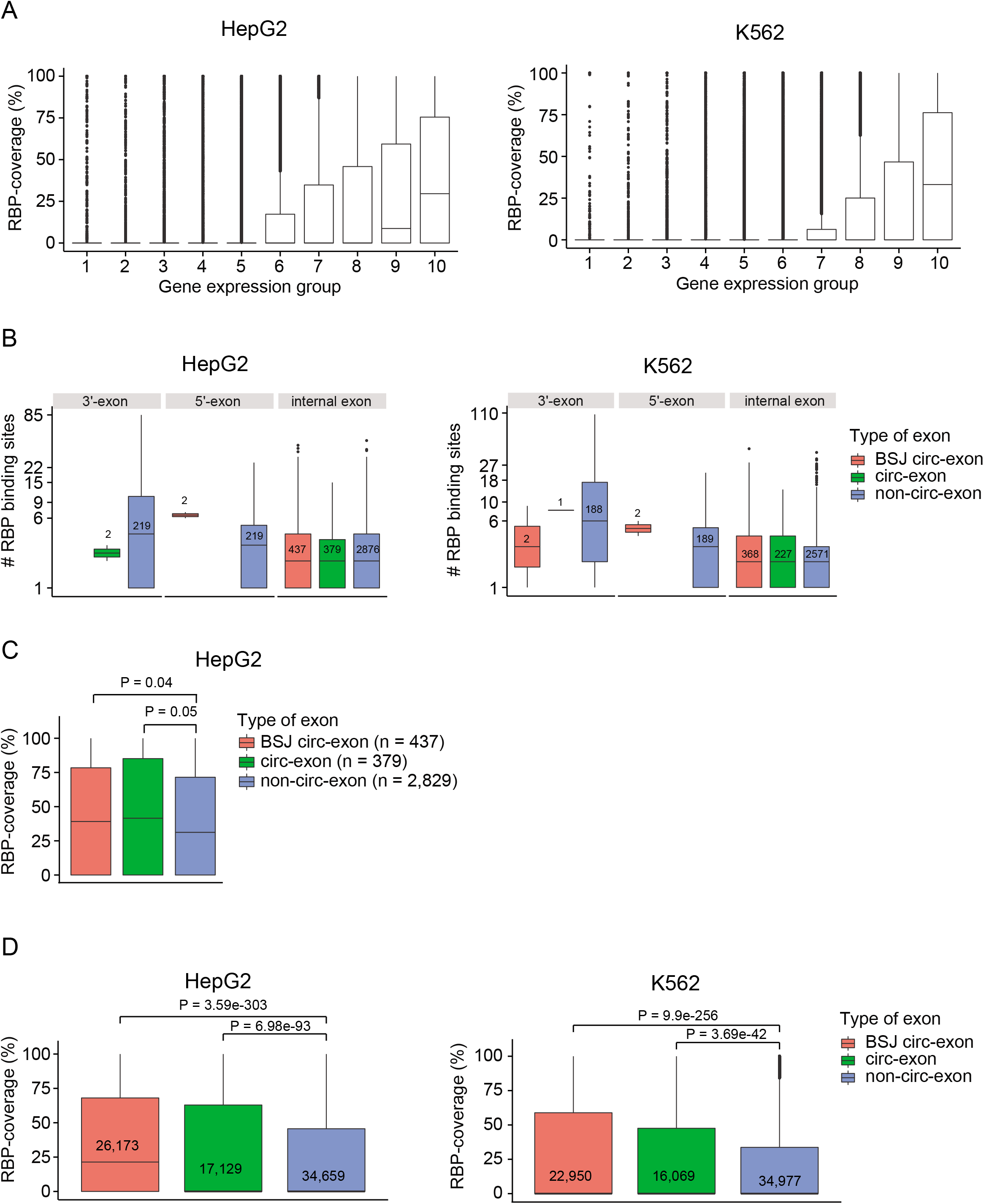
Exons comprising circRNAs are enriched with RBP binding sites 3A) Fraction of exons covered with RBP binding sites in different gene expression groups. All expressed genes in HepG2 (n = 18,152) and K562 (n = 18,784) were divided into ten bins based on expression. RBP-coverage correlate with gene expression in HepG2 (r = 0.41, P < 2.2e-16, Pearson’s product-moment correlation) and K562 (r = 0.42, P < 2.2e-16, Pearson’s product-moment correlation). **3B)** Number of RBP binding sites in 5’-, 3’-, and internal exons. Numbers represent number of exons in each group. The y-axis is plotted on a log10-scale but showing real numbers of RBP binding sites. **3C)** RBP binding site coverage for each category of exons in genes that produce highly expressed (top 1%) circRNAs in HepG2 (n = 218). P-values obtained by Wilcoxon Rank Sum Test. **3D)** RBP-coverage for each category of exons in genes that produce circRNAs (supported by at least two reads) in HepG2 (n = 5,284 genes) and K562 (n = 5,244 genes). Numbers represent number of exons in each group. P-values obtained by Wilcoxon Rank Sum Test. **3E)** Gene expression group distribution of genes producing highly expressed circRNAs in HepG2 (n = 218 genes) and K562 (n = 185). In cases where more than one circRNA arise from the same gene, the gene is only counted once. **3F)** Comparison of RBP-coverage between BSJ circ-exons and exons in groups of genes of different expression levels (HepG2). The mean RBP-coverage of the highly expressed BSJ circ-exons (n = 437) is 42% (red punctuated line; median = 40%). Exons randomly sampled from genes while ensuring the same expression profile as genes producing highly expressed circRNAs (circ-genes) have a much lower mean coverage of 25% (green; median = 0%). Exons of highly expressed genes (top 5-10 percentiles) also showed a lower mean coverage of 33% (purple; median = 13%), while the most highly expressed genes (top 5 percentiles) had a mean of 39% (blue; median = 30%). The random sampling procedures were repeated with 100 iterations. Empirical P-values for: BSJ circ-exons vs. same expression, P < 0.01; BSJ circ-exons vs. Top5-10, P < 0.01; BSJ circ-exons vs Top5, P = 0.1. **3G)** Mean RBP-coverage per exon for individual RBPs in HepG2. All RBPs shown here have significantly more target sites in BSJ circ-exons than non-circ-exons of the same genes (FDR < 0.1, Wilcoxon Rank Sum Test). Only RBPs with at least 20 distinct binding sites in total are considered. **3H)** Approach for identifying eCLIP reads spanning the backsplice junction (BSJ) of circRNAs. By extracting 30 bp from each side of all exons, we generated a reference set of all possible exonic BSJ events. We extracted all unmapped eCLIP reads and mapped them against the BSJ reference set, allowing two mismatches. Next, we removed PCR duplicates and merged barcode replicates. We removed read 1 (which was only used to identify and remove PCR duplicates) and counted eCLIP reads spanning each BSJ. Only reads spanning a BSJ by at least 5 bp were counted. To identify specific RBP binding sites, we only reported binding sites supported by at least 10 reads in both IP replicates and with an 8-fold enrichment in IP compared to input (Methods). **3I)** Number of eCLIP reads spanning BSJ events in HepG2 (left) and K562 (right). Some BSJ events are supported by eCLIP reads, however, none of these correspond to a circRNA predicted by CIRI2.

**Supplementary Figure 4:**
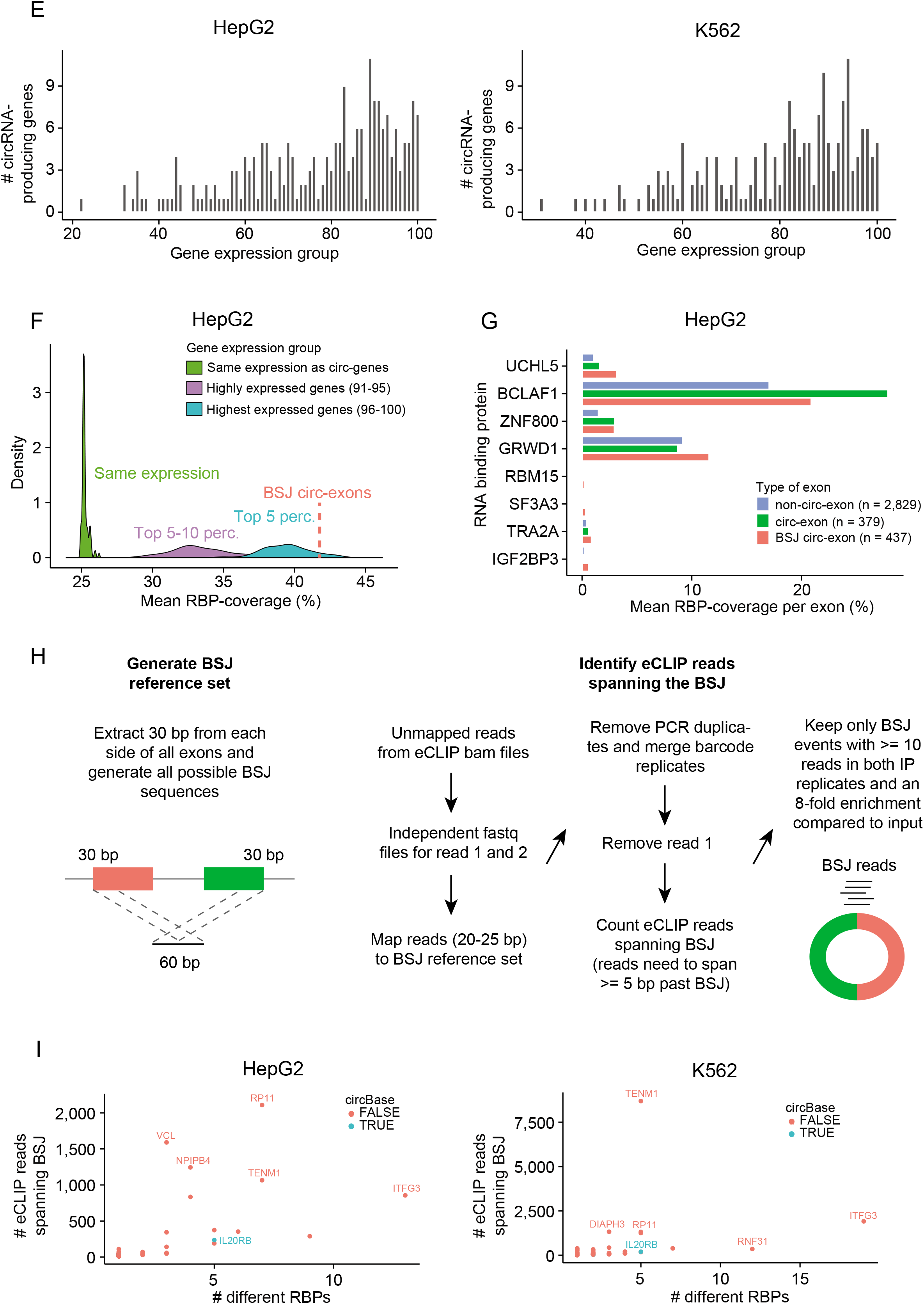
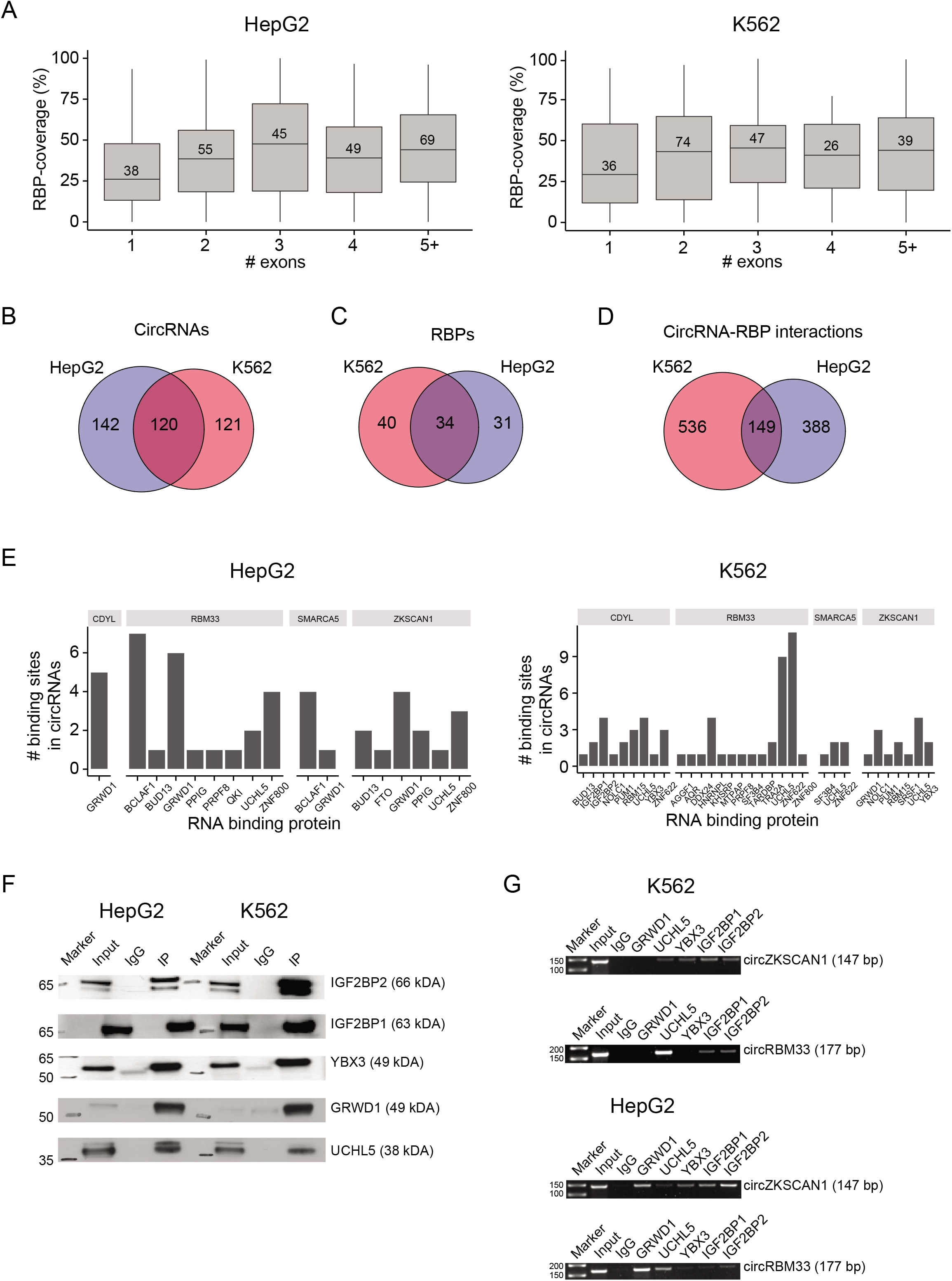
CircRNAs interact with RBPs in a cell-type-specific manner. **4A)** RBP-coverage based on number of exons comprising the circRNAs in HepG2 (left) and K562 (right). Numbers indicate the number of circRNAs belonging to each group. **4B)** Of the 1% highest expressed circRNAs in HepG2 (n = 262) and K562 (n = 241), 120 circRNAs are highly expressed in both cell lines. **4C)** Of RBPs with binding sites in the shared highly expressed circRNAs in HepG2 (n = 65) and K562 (n = 74), 34 RBPs are evaluated in both cell lines. **4D)** CircRNA-RBP interactions identified in both cell lines. 149 interactions between 120 highly expressed circRNAs and 34 RBPs are identified in both HepG2 and K562. The genomic positions of RBP binding sites are not considered. **4E)** Number of predicted binding sites for each RBP in four highly expressed and RBP-covered circRNAs; circCDYL, circRBM33, circSMARCA5, and circZKSCAN1. **4F)** Western blot of IP and input from the RIP experiments. IgG is used as negative control. **4G)** Validation of circRNA-RBP interactions from RIP experiments. Here, RIP experiments confirmed that circRBM33 interacts strongly with UCHL5 in K562 and with both GRWD1 and UCHL5 in HepG2. For circZKSCAN1, several interactions were observed, however, binding affinities were generally low. Specifically, circZKSCAN1 contain a single binding site for GRWD1 in K562 (Supp. Figure 4E), but no interaction was detected from the RIP experiments. 50 bp markers used.

**Supplementary Figure 5:**
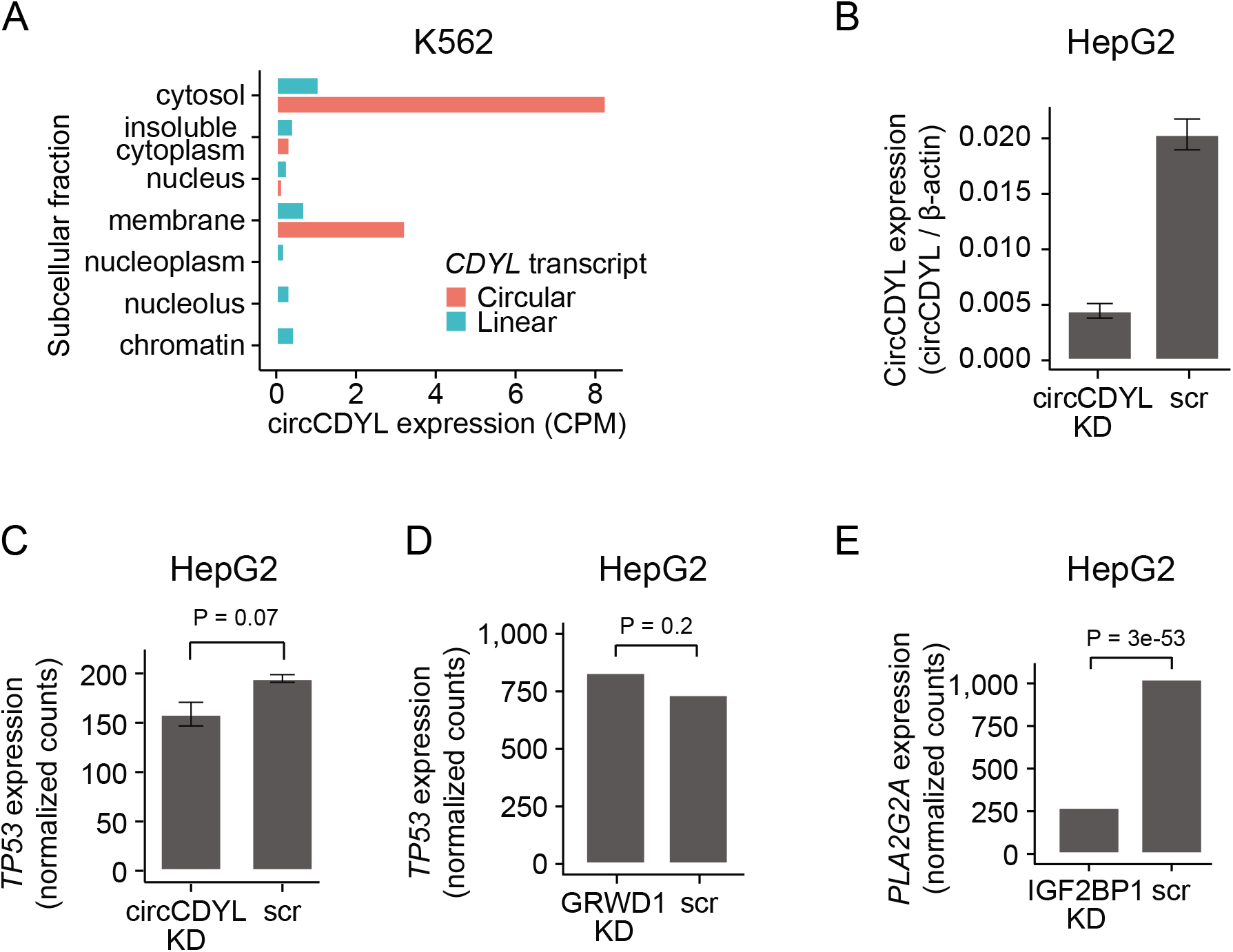
Functional studies of circCDYL-GRWD1 interactions in HepG2. **5A)** Expression of circCDYL and the corresponding linear transcript in subcellular fractions of K562. **5B)** Knockdown efficiency of circCDYL in HepG2 cells. **5C+D)** Expression of *TP53* upon circCDYL KD (C) and GRWD1 KD (D) in HepG2 cells. P-values obtained by Wald Test. **5E)** Expression of *PLA2G2A* upon IGF2BP1 KD in HepG2 cells. P-value obtained by Wald Test.

**Supplementary Figure 6:**
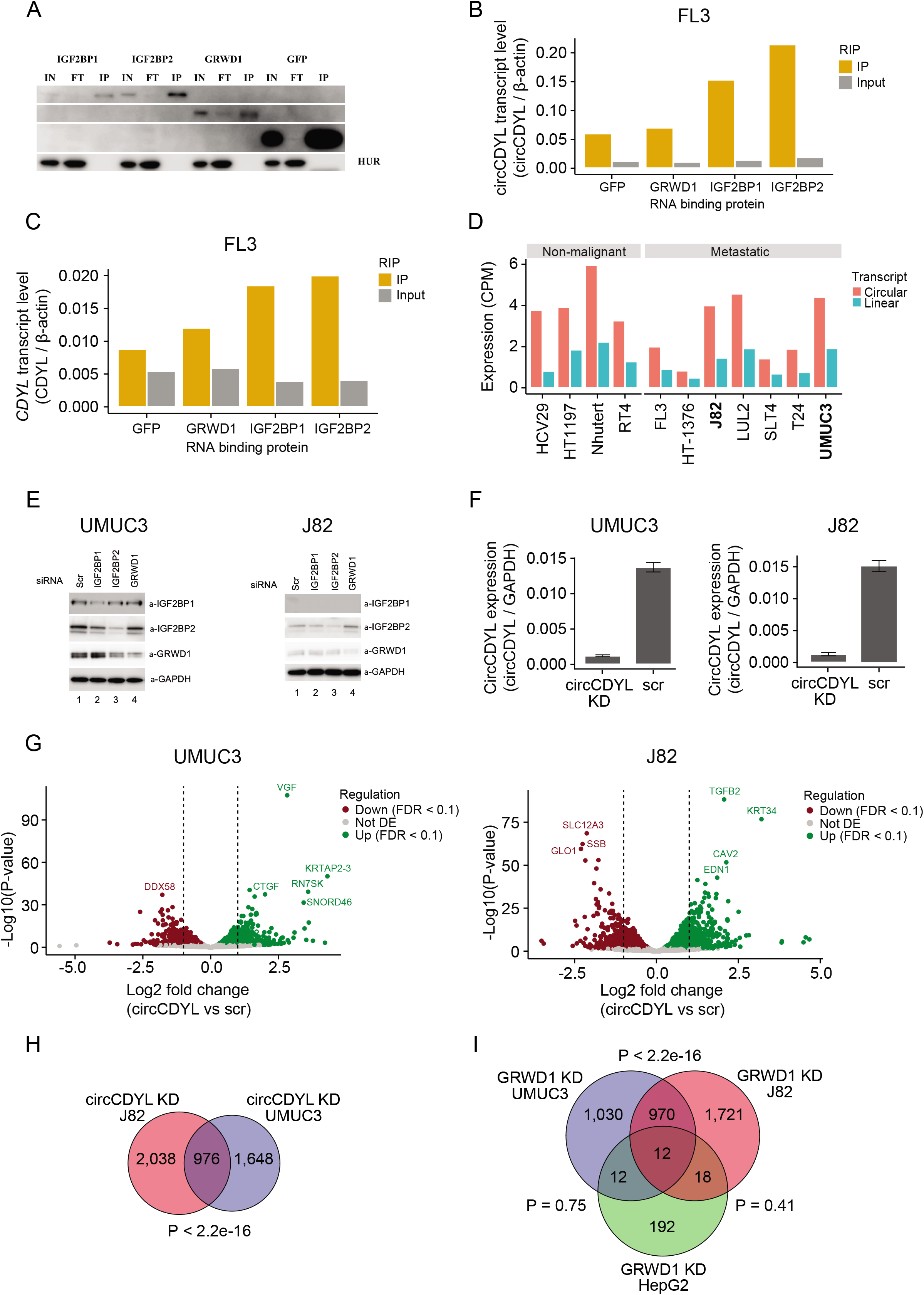
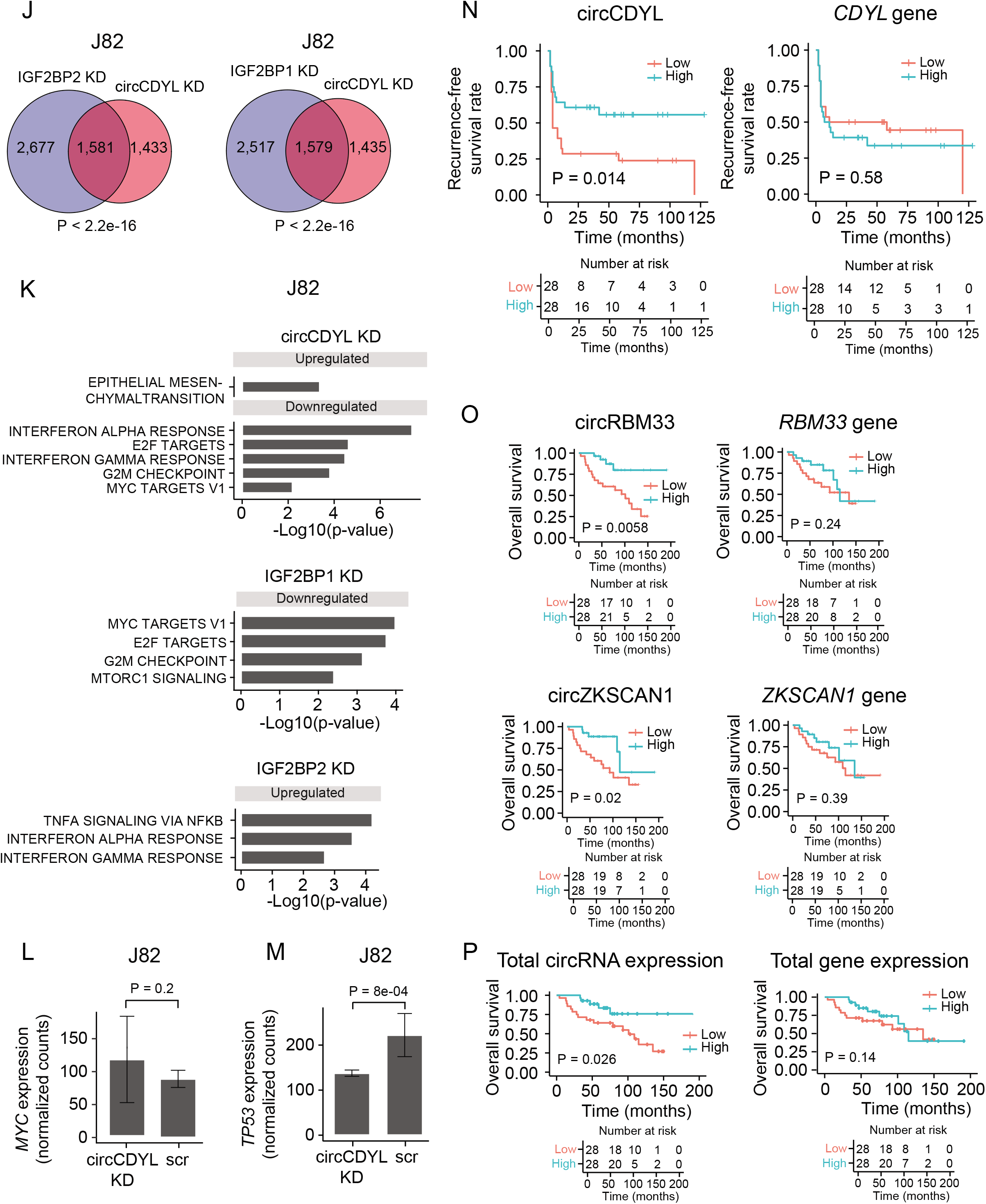
CircCDYL interacts with RBPs in bladder cancer and abundance is associated with overall survival. **6A)** Validation of IP by western blot. Overexpression of Twin-Strep-tagged RBPs for 48 hours followed by IP and RNA purification. Experiments are performed in the bladder cancer cell line FL3. IN = Input, FT = Flow through, IP = IP of RBP. **6B+C)** CircCDYL (B) and CDYL (C) transcript expression levels relative to β-actin expression levels for IP and input in FL3. GFP was used as control. **6D)** Expression of circCDYL and the corresponding linear transcript in eleven bladder cancer cell lines. UMUC3 and J82 were chosen for knockdown experiments based on circCDYL expression levels and cell line stability. **6E)** Western blots of IGF2BP1, IGF2BP2, and GRWD1 knockdown in UMUC3 and J82 cells. **6F)** Knockdown efficiency of circCDYL in UMUC3 and J82 cells. **6G)** Differential expression analyses of mRNAs upon circCDYL KD in J82 (top) and UMUC3 (bottom). The log2 fold changes (circCDYL KD vs scr) are plotted against the negative log10(p-values). Colors indicate if genes are significantly down-(red) or upregulated (green) or not significantly differentially expressed (Not DE, grey) after Benjamini-Hochberg correction, FDR < 0.1. Vertical lines indicate a log2FC > 1 or < −1.**6H)** Overlap of differentially expressed genes upon circCDYL KD in J82 and UMUC3 cells. P-value obtained by Fisher’s Exact Test. **6I)** Overlap of genes affected by GRWD1 KD in HepG2, UMUC3, and J82 cells. P-values obtained by Fisher’s Exact Test. **6J)** Overlap of genes affected by circCDYL KD, and IGF2BP1 and IGF2BP2, respectively, in J82. P-values obtained by Fisher’s Exact Test. **6K)** Gene set enrichment analysis of 50 hallmarks of cancer upon circCDYL KD (top), IGF2BP1 KD (middle), and IGF2BP2 KD (bottom) in J82 cells. **6L+M)** Expression of *MYC* (K) and *TP53* (L) upon circCDYL KD in J82. P-values obtained by Wald Test. **6N)** Kaplan-Meier recurrence-free survival plots for circCDYL and *CDYL* host gene. Median expression used as cutoff; circCDYL = 0.125 CPM and *CDYL* = 21.4 FPKM. P-values obtained by Log-Rank Test. **6O)** Kaplan-Meier overall survival plots for circRBM33, *RBM33* host gene, circZKSCAN1, and *ZKSCAN1* host gene. Median expression used as cutoff; circRBM33 = 0.0618 CPM, *RBM33* = 21.3 FPKM, circZKSCAN1 = 0.0994 CPM, and *ZKSCAN1* = 17.8 FPKM. P-values obtained by Log-Rank Test. **6P)** Kaplan-Meier overall survival plots for total circRNA and gene expression. Median expression used as cutoff; circRNAs = 19.5 CPM and genes = 366,000 FPKM. P-values obtained by Log-Rank Test.

## References

1. Starke, S., Jost, I., Rossbach, O., Schneider, T., Schreiner, S., Hung, L.-H. and Bindereif, A. (2015) Exon Circularization Requires Canonical Splice Signals. Cell Reports, 10, 103–111.

2. Cocquerelle, C., Mascrez, B., Hétuin, D. and Bailleul, B. (1993) Mis-splicing yields circular RNA molecules. FASEB J., 7, 155–160.

3. Memczak, S., Jens, M., Elefsinioti, A., Torti, F., Krueger, J., Rybak, A., Maier, L., Mackowiak, S.D., Gregersen, L.H., Munschauer, M., et al. (2013) Circular RNAs are a large class of animal RNAs with regulatory potency. Nature, 495, 333–338.

4. Jeck, W.R., Sorrentino, J.A., Wang, K., Slevin, M.K., Burd, C.E., Liu, J., Marzluff, W.F. and Sharpless, N.E. (2013) Circular RNAs are abundant, conserved, and associated with ALU repeats. RNA, 19, 141–157.

5. Salzman, J., Gawad, C., Wang, P.L., Lacayo, N. and Brown, P.O. (2012) Circular RNAs are the predominant transcript isoform from hundreds of human genes in diverse cell types. PLoS One, 7, e30733.

6. Zhang, Z., Yang, T. and Xiao, J. (2018) Circular RNAs: Promising Biomarkers for Human Diseases. EBioMedicine, 34, 267–274.

7. Chen, S., Huang, V., Xu, X., Livingstone, J., Soares, F., Jeon, J., Zeng, Y., Hua, J.T., Petricca, J., Guo, H., et al. (2019) Widespread and Functional RNA Circularization in Localized Prostate Cancer. Cell, 176, 831–843. e22.

8. Vo, J.N., Cieslik, M., Zhang, Y., Shukla, S., Xiao, L., Zhang, Y., Wu, Y.-M., Dhanasekaran, S.M., Engelke, C.G., Cao, X., et al. (2019) The Landscape of Circular RNA in Cancer. Cell, 176, 869–881. e13.

9. Li, Y., Zheng, Q., Bao, C., Li, S., Guo, W., Zhao, J., Chen, D., Gu, J., He, X. and Huang, S. (2015) Circular RNA is enriched and stable in exosomes: a promising biomarker for cancer diagnosis. Cell Res., 25, 981–984.

10. Memczak, S., Papavasileiou, P., Peters, O. and Rajewsky, N. (2015) Identification and Characterization of Circular RNAs As a New Class of Putative Biomarkers in Human Blood. PLoS One, 10, e0141214.

11. Bahn, J.H., Zhang, Q., Li, F., -M. Chan, T., Lin, X., Kim, Y., Wong, D.T.W. and Xiao, X. (2015) The Landscape of MicroRNA, Piwi-Interacting RNA, and Circular RNA in Human Saliva. Clinical Chemistry, 61, 221–230.

12. Dominguez, D., Freese, P., Alexis, M.S., Su, A., Hochman, M., Palden, T., Bazile, C., Lambert, N.J., Van Nostrand, E.L., Pratt, G.A., et al. (2018) Sequence, Structure, and Context Preferences of Human RNA Binding Proteins. Mol. Cell, 70, 854–867. e9.

13. Glisovic, T., Bachorik, J.L., Yong, J. and Dreyfuss, G. (2008) RNA-binding proteins and post-transcriptional gene regulation. FEBS Letters, 582, 1977–1986.

14. Zang, J., Lu, D. and Xu, A. (2018) The interaction of circRNAs and RNA binding proteins: An important part of circRNA maintenance and function. J. Neurosci. Res., 10.1002/jnr.24356.

15. Conn, S.J., Pillman, K.A., Toubia, J., Conn, V.M., Salmanidis, M., Phillips, C.A., Roslan, S., Schreiber, A.W., Gregory, P.A. and Goodall, G.J. (2015) The RNA binding protein quaking regulates formation of circRNAs. Cell, 160, 1125–1134.

16. Errichelli, L., Dini Modigliani, S., Laneve, P., Colantoni, A., Legnini, I., Capauto, D., Rosa, A., De Santis, R., Scarfò, R., Peruzzi, G., et al. (2017) FUS affects circular RNA expression in murine embryonic stem cell-derived motor neurons. Nat. Commun., 8, 14741.

17. Fei, T., Chen, Y., Xiao, T., Li, W., Cato, L., Zhang, P., Cotter, M.B., Bowden, M., Lis, R.T., Zhao, S.G., et al. (2017) Genome-wide CRISPR screen identifies HNRNPL as a prostate cancer dependency regulating RNA splicing. Proceedings of the National Academy of Sciences, 10.1073/pnas.1617467114.

18. Khan, M.A.F., Reckman, Y.J., Aufiero, S., van den Hoogenhof, M.M.G., van der Made, I., Beqqali, A., Koolbergen, D.R., Rasmussen, T.B., van der Velden, J., Creemers, E.E., et al. (2016) RBM20 Regulates Circular RNA Production From the Titin Gene. Circ. Res, 119, 996–1003.

19. Ashwal-Fluss, R., Meyer, M., Pamudurti, N.R., Ivanov, A., Bartok, O., Hanan, M., Evantal, N., Memczak, S., Rajewsky, N. and Kadener, S. (2014) circRNA biogenesis competes with pre-mRNA splicing. Mol. Cell, 56, 55–66.

20. Li, X., Liu, C.-X., Xue, W., Zhang, Y., Jiang, S., Yin, Q.-F., Wei, J., Yao, R.-W., Yang, L. and Chen, L.-L. (2017) Coordinated circRNA Biogenesis and Function with NF90/NF110 in Viral Infection. Mol. Cell, 67, 214–227. e7.

21. Ivanov, A., Memczak, S., Wyler, E., Torti, F., Porath, H.T., Orejuela, M.R., Piechotta, M., Levanon, E.Y., Landthaler, M., Dieterich, C., et al. (2015) Analysis of intron sequences reveals hallmarks of circular RNA biogenesis in animals. Cell Rep, 10, 170–177.

22. Aktaş, T., Avşar Ilik, I., Maticzka, D., Bhardwaj, V., Pessoa Rodrigues, C., Mittler, G., Manke, T., Backofen, R. and Akhtar, A. (2017) DHX9 suppresses RNA processing defects originating from the Alu invasion of the human genome. Nature, 544, 115–119.

23. Hansen, T.B., Jensen, T.I., Clausen, B.H., Bramsen, J.B., Finsen, B., Damgaard, C.K. and Kjems, J. (2013) Natural RNA circles function as efficient microRNA sponges. Nature, 495, 384–388.

24. Piwecka, M., Glažar, P., Hernandez-Miranda, L.R., Memczak, S., Wolf, S.A., Rybak-Wolf, A., Filipchyk, A., Klironomos, F., Cerda Jara, C.A., Fenske, P., et al. (2017) Loss of a mammalian circular RNA locus causes miRNA deregulation and affects brain function. Science, 357.

25. Kleaveland, B., Shi, C.Y., Stefano, J. and Bartel, D.P. (2018) A Network of Noncoding Regulatory RNAs Acts in the Mammalian Brain. Cell, 174, 350–362. e17.

26. Abdelmohsen, K., Panda, A.C., Munk, R., Grammatikakis, I., Dudekula, D.B., De, S., Kim, J., Noh, J.H., Kim, K.M., Martindale, J.L., et al. (2017) Identification of HuR target circular RNAs uncovers suppression of PABPN1 translation by CircPABPN1. RNA Biol., 14, 361–369.

27. Du, W.W., Yang, W., Liu, E., Yang, Z., Dhaliwal, P. and Yang, B.B. (2016) Foxo3 circular RNA retards cell cycle progression via forming ternary complexes with p21 and CDK2. Nucleic Acids Res., 44, 2846–2858.

28. Du, W.W., Yang, W., Chen, Y., Wu, Z.-K., Foster, F.S., Yang, Z., Li, X. and Yang, B.B. (2017) Foxo3 circular RNA promotes cardiac senescence by modulating multiple factors associated with stress and senescence responses. Eur. Heart J., 38, 1402–1412.

29. Schneider, T., Hung, L.-H., Schreiner, S., Starke, S., Eckhof, H., Rossbach, O., Reich, S., Medenbach, J. and Bindereif, A. (2016) CircRNA-protein complexes: IMP3 protein component defines subfamily of circRNPs. Sci. Rep., 6, 31313.

30. You, X., Vlatkovic, I., Babic, A., Will, T., Epstein, I., Tushev, G., Akbalik, G., Wang, M., Glock, C., Quedenau, C., et al. (2015) Neural circular RNAs are derived from synaptic genes and regulated by development and plasticity. Nat. Neurosci., 18, 603–610.

31. Hong, S. (2017) RNA Binding Protein as an Emerging Therapeutic Target for Cancer Prevention and Treatment. J Cancer Prev, 22, 203–210.

32. Van Nostrand, E.L., Pratt, G.A., Shishkin, A.A., Gelboin-Burkhart, C., Fang, M.Y., Sundararaman, B., Blue, S.M., Nguyen, T.B., Surka, C., Elkins, K., et al. (2016) Robust transcriptome-wide discovery of RNA-binding protein binding sites with enhanced CLIP (eCLIP). Nat. Methods, 13, 508–514.

33. ENCODE Project Consortium (2012) An integrated encyclopedia of DNA elements in the human genome. Nature, 489, 57–74.

34. Gao, Y., Zhang, J. and Zhao, F. (2018) Circular RNA identification based on multiple seed matching. Brief. Bioinform., 19, 803–810.

35. Glažar, P., Papavasileiou, P. and Rajewsky, N. (2014) circBase: a database for circular RNAs. RNA, 20, 1666–1670.

36. Kristensen, L.S., Okholm, T.L.H., Venø, M.T. and Kjems, J. (2018) Circular RNAs are abundantly expressed and upregulated during human epidermal stem cell differentiation. RNA Biol., 15, 280.

37. Wahl, M.C., Will, C.L. and Lührmann, R. (2009) The spliceosome: design principles of a dynamic RNP machine. Cell, 136, 701–718.

38. Van Nostrand, E.L., Freese, P., Pratt, G.A., Wang, X., Wei, X., Xiao, R., Blue, S.M., Chen, J.-Y., Cody, N.A.L., Dominguez, D., et al. (2018) A Large-Scale Binding and Functional Map of Human RNA Binding Proteins. bioRxiv, 10.1101/179648.

39. Zhang, J., Zhang, X., Li, C., Yue, L., Ding, N., Riordan, T., Yang, L., Li, Y., Jen, C., Lin, S., et al. (2019) Circular RNA profiling provides insights into their subcellular distribution and molecular characteristics in HepG2 cells. RNA Biol., 16, 220–232.

40. Briata, P., Bordo, D., Puppo, M., Gorlero, F., Rossi, M., Bizzozzero, N.P. and Gherzi, R. (2016) Diverse roles of the nucleic acid binding protein KHSRP in cell differentiation and disease. Wiley Interdiscip. Rev. RNA, 7, 227.

41. Min, H., Turck, C.W., Nikolic, J.M. and Black, D.L. (1997) A new regulatory protein, KSRP, mediates exon inclusion through an intronic splicing enhancer. Genes Dev., 11, 1023–1036.

42. Singh, G., Pratt, G., Yeo, G.W. and Moore, M.J. (2015) The Clothes Make the mRNA: Past and Present Trends in mRNP Fashion. Annu. Rev. Biochem., 84, 325–354.

43. Stagsted, L.V., Nielsen, K.M., Daugaard, I. and Hansen, T.B. (2019) Noncoding AUG circRNAs constitute an abundant and conserved subclass of circles. Life Sci Alliance, 2.

44. Voellenkle, C., Perfetti, A., Carrara, M., Fuschi, P., Renna, L.V., Longo, M., Sain, S.B., Cardani, R., Valaperta, R., Silvestri, G., et al. (2019) Dysregulation of Circular RNAs in Myotonic Dystrophy Type 1. Int.J. Mol. Sci., 20.

45. Okholm, T.L.H., Nielsen, M.M., Hamilton, M.P., Christensen, L.-L., Vang, S., Hedegaard, J., Hansen, T.B., Kjems, J., Dyrskjøt, L. and Pedersen, J.S. (2017) Circular RNA expression is abundant and correlated to aggressiveness in early-stage bladder cancer. npj Genomic Medicine, 2.

46. Moll, P., Ante, M., Seitz, A. and Reda, T. (2014) QuantSeq 3′ mRNA sequencing for RNA quantification. Nature Methods, 11, i–iii.

47. Takafuji, T., Kayama, K., Sugimoto, N. and Fujita, M. (2017) GRWD1, a new player among oncogenesis-related ribosomal/nucleolar proteins. Cell Cycle, 16, 1397.

48. Sugimoto, N., Maehara, K., Yoshida, K., Yasukouchi, S., Osano, S., Watanabe, S., Aizawa, M., Yugawa, T., Kiyono, T., Kurumizaka, H., et al. (2015) Cdt1-binding protein GRWD1 is a novel histone-binding protein that facilitates MCM loading through its influence on chromatin architecture. Nucleic Acids Res., 43, 5898–5911.

49. Kayama, K., Watanabe, S., Takafuji, T., Tsuji, T., Hironaka, K., Matsumoto, M., Nakayama, K.I., Enari, M., Kohno, T., Shiraishi, K., et al. (2017) GRWD1 negatively regulates p53 via the RPL11-MDM2 pathway and promotes tumorigenesis. EMBO Rep., 18, 123–137.

50. Huang, X., Zhang, H., Guo, X., Zhu, Z., Cai, H. and Kong, X. (2018) Insulin-like growth factor 2 mRNA-binding protein 1 (IGF2BP1) in cancer. J. Hematol. Oncol., 11, 88.

51. Stöhr, N., Köhn, M., Lederer, M., Glass, M., Reinke, C., Singer, R.H. and Hüttelmaier, S. (2012) IGF2BP1 promotes cell migration by regulating MK5 and PTEN signaling. Genes Dev., 26, 176–189.

52. Rini, J. and Anbalagan, M. (2017) IGF2BP1: a novel binding protein of p38 MAPK. Mol. Cell. Biochem., 435, 133–140.

53. Xu, Y., Zheng, Y., Liu, H. and Li, T. (2017) Modulation of IGF2BP1 by long non-coding RNA HCG11 suppresses apoptosis of hepatocellular carcinoma cells via MAPK signaling transduction. Int. J. Oncol., 51, 791–800.

54. Kennedy, B.P., Soravia, C., Moffat, J., Xia, L., Hiruki, T., Collins, S., Gallinger, S. and Bapat, B. (1998) Overexpression of the nonpancreatic secretory group II PLA2 messenger RNA and protein in colorectal adenomas from familial adenomatous polyposis patients. Cancer Res., 58, 500–503.

55. Buhmeida, A., Bendardaf, R., Hilska, M., Laine, J., Collan, Y., Laato, M., Syrjänen, K. and Pyrhönen, S. (2009) PLA2 (group IIA phospholipase A2) as a prognostic determinant in stage II colorectal carcinoma. Ann. Oncol., 20, 1230–1235.

56. He, H.-L., Lee, Y.-E., Shiue, Y.-L., Lee, S.-W., Lin, L.-C., Chen, T.-J., Wu, T.-F. and Li, C.-F. (2015) PLA2G2A overexpression is associated with poor therapeutic response and inferior outcome in rectal cancer patients receiving neoadjuvant concurrent chemoradiotherapy. Histopathology, 66, 991–1002.

57. Hedegaard, J., Lamy, P., Nordentoft, I., Algaba, F., Høyer, S., Ulhøi, B.P., Vang, S., Reinert, T., Hermann, G.G., Mogensen, K., et al. (2016) Comprehensive Transcriptional Analysis of Early-Stage Urothelial Carcinoma. Cancer Cell, 30, 27–42.

58. Sun, J., Zhang, H., Tao, D., Xie, F., Liu, F., Gu, C., Wang, M., Wang, L., Jiang, G., Wang, Z., et al. (2019) CircCDYL inhibits the expression of C-MYC to suppress cell growth and migration in bladder cancer. Artif. Cells Nanomed. Biotechnol., 47, 1349–1356.

59. Cao, J., Mu, Q. and Huang, H. (2018) The Roles of Insulin-Like Growth Factor 2 mRNA-Binding Protein 2 in Cancer and Cancer Stem Cells. Stem Cells Int., 2018, 4217259.

60. Liberzon, A., Birger, C., Thorvaldsdóttir, H., Ghandi, M., Mesirov, J.P. and Tamayo, P. (2015) The Molecular Signatures Database Hallmark Gene Set Collection. Cell Systems, 1, 417–425.

61. Tate, J.G., Bamford, S., Jubb, H.C., Sondka, Z., Beare, D.M., Bindal, N., Boutselakis, H., Cole, C.G., Creatore, C., Dawson, E., et al. (2019) COSMIC: the Catalogue Of Somatic Mutations In Cancer. Nucleic Acids Res., 47, D941–D947.

62. Jakobi, T. and Dieterich, C. (2019) Computational approaches for circular RNA analysis. Wiley Interdisciplinary Reviews: RNA, 10, e1528.

63. Jeck, W.R. and Sharpless, N.E. (2014) Detecting and characterizing circular RNAs. Nat. Biotechnol., 32, 453–461.

64. Hansen, T.B., Kjems, J. and Damgaard, C.K. (2013) Circular RNA and miR-7 in Cancer. Cancer Research, 73, 5609–5612.

65. Chen, Y.G., Kim, M.V., Chen, X., Batista, P.J., Aoyama, S., Wilusz, J.E., Iwasaki, A. and Chang, H.Y. (2017) Sensing Self and Foreign Circular RNAs by Intron Identity. Mol. Cell, 67, 228–238. e5.

66. Du, W.W., Fang, L., Yang, W., Wu, N., Awan, F.M., Yang, Z. and Yang, B.B. (2017) Induction of tumor apoptosis through a circular RNA enhancing Foxo3 activity. Cell Death Differ., 24, 357–370.

67. Du, W.W., Zhang, C., Yang, W., Yong, T., Awan, F.M. and Yang, B.B. (2017) Identifying and Characterizing circRNA-Protein Interaction. Theranostics, 7, 4183–4191.

68. Kim, D., Pertea, G., Trapnell, C., Pimentel, H., Kelley, R. and Salzberg, S.L. (2013) TopHat2: accurate alignment of transcriptomes in the presence of insertions, deletions and gene fusions. Genome Biol., 14, R36.

69. Langmead, B. and Salzberg, S.L. (2012) Fast gapped-read alignment with Bowtie 2. Nat. Methods, 9, 357–359.

70. Trapnell, C., Williams, B.A., Pertea, G., Mortazavi, A., Kwan, G., Van Baren, M.J., Salzberg, S.L., Wold, B.J. and Pachter, L. (2010) Transcript assembly and quantification by RNA-Seq reveals unannotated transcripts and isoform switching during cell differentiation. Nat. Biotechnol., 28, 511–515.

71. Anders, S., Pyl, P.T. and Huber, W. (2015) HTSeq--a Python framework to work with high-throughput sequencing data. Bioinformatics, 31, 166–169.

72. Li, H., Handsaker, B., Wysoker, A., Fennell, T., Ruan, J., Homer, N., Marth, G., Abecasis, G., Durbin, R. and 1000 Genome Project Data Processing Subgroup (2009) The Sequence Alignment/Map format and SAMtools. Bioinformatics, 25, 2078–2079.

73. Lovci, M.T., Ghanem, D., Marr, H., Arnold, J., Gee, S., Parra, M., Liang, T.Y., Stark, T.J., Gehman, L.T., Hoon, S., et al. (2013) Rbfox proteins regulate alternative mRNA splicing through evolutionarily conserved RNA bridges. Nat. Struct. Mol. Biol., 20, 1434–1442.

74. Zerbino, D.R., Achuthan, P., Akanni, W., Amode, M.R., Barrell, D., Bhai, J., Billis, K., Cummins, C., Gall, A., Girón, C.G., et al. (2018) Ensembl 2018. Nucleic Acids Res., 46, D754–D761.

75. Luo, W., Friedman, M.S., Shedden, K., Hankenson, K.D. and Woolf, P.J. (2009) GAGE: generally applicable gene set enrichment for pathway analysis. BMC Bioinformatics, 10, 161.

76. Thomsen, R., Sølvsten, C.A.E., Linnet, T.E., Blechingberg, J. and Nielsen, A.L. (2010) Analysis of qPCR data by converting exponentially related Ct values into linearly related X0 values. J. Bioinform. Comput. Biol., 8, 885–900.

77. R Core Team (2019) R: A language and environment for statistical computing. R Foundation for Statistical Computing, Vienna, Austria.

78. RStudio Team (2016) RStudio: Integrated Development for R. RStudio, Inc., Boston, MA.

79. Love, M.I., Huber, W. and Anders, S. (2014) Moderated estimation of fold change and dispersion for RNA-seq data with DESeq2. Genome Biol., 15, 550.

80. Therneau, T. (2015) A Package for Survival Analysis in S. Version 2.38.

81. Therneau, T.M. and Grambsch, P.M. (2000) Modeling Survival Data: Extending the Cox Model. Springer, New York.

82. Kassambara, A., Kosinski, M. and Biecek, P. (2019) survminer: Drawing Survival Curves using ‘ggplot2’. R package version 0.4.6.

83. Wickham, H. (2016) ggplot2: Elegant Graphics for Data Analysis Springer.

